# The *Dlk1-Dio3* noncoding RNA cluster coordinately regulates mitochondrial respiration and chromatin structure to establish proper cell state for muscle differentiation

**DOI:** 10.1101/2024.06.18.599546

**Authors:** Amanda Pinheiro, Christopher A. Petty, Kevin Cabrera, Eric P. Tost, Adam C. Gower, Madison Marano, Ethan M. Leviss, Matthew J. Boberg, Jawahar Mahendran, Payton M. Bock, Chelsea E. Stephens, Jessica L. Fetterman, Francisco J. Naya

## Abstract

The coordinate regulation of metabolism and epigenetics to establish cell state-specific gene expression patterns during lineage progression is a central aspect of cell differentiation, but the factors that regulate this elaborate interplay are not well-defined. The imprinted *Dlk1-Dio3* noncoding RNA (ncRNA) cluster has been associated with metabolism in various progenitor cells, suggesting it functions as a regulator of metabolism and cell state. Here, we directly demonstrate that the *Dlk1-Dio3* ncRNA cluster coordinates mitochondrial respiration and chromatin structure to maintain proper cell state. Stable muscle cell lines were generated harboring two distinct deletions in the proximal promoter region resulting in either greatly upregulated or downregulated expression of the entire *Dlk1-Dio3* ncRNA cluster. Both mutant lines displayed impaired muscle differentiation along with altered mitochondrial respiration and genome-wide changes in chromatin accessibility and histone methylation. Global gene expression patterns and pathway analyses indicated a reprogramming of myogenic cell state creating a differentiated-like phenotype in proliferating myoblasts. Our results strongly suggest the *Dlk1-Dio3* ncRNA locus is a nodal regulator coordinating metabolic activity and the epigenome to maintain proper cell state in the myogenic lineage.

**Summary statement:** Muscle cell state is regulated by the imprinted *Dlk1-Dio3* noncoding RNA locus through its coordinate control of mitochondrial activity and histone modifications.

## Introduction

Crosstalk between metabolic pathways and the epigenome is an essential node in the regulatory network of cell state transitions in development and disease (Chakraborty and Chandel 2021; Folmes, et al. 2012; Ly et al., 2020; Tarazona et al. 2020). A central aspect of cell state involves the establishment of genome-wide histone modifications that induce changes in chromatin structure to alter gene expression patterns characteristic of and necessary for a given cell state. These cell state-specific epigenetic landscapes are typically mediated by the concerted function of chromatin-modifying enzymes and transcription factors. Many chemical reactions utilized for the modification of histones require intermediate metabolites produced by metabolic pathways (Etchegaray and Mostoslavsky 2016; Reid et al., 2017). Consequently, dynamic changes in histone modifications are known to parallel changes in cellular metabolism.

Muscle differentiation serves as an excellent paradigm to investigate the interplay between metabolism and cell state. As muscle progenitors differentiate, each transitory cell state has unique energy demands associated with a predominant energy-producing metabolic pathway (Bhattacharya and Scime, 2020; Relaix et al., 2021). Proliferating myoblasts, for example, utilize glycolysis as their optimal energy source to support robust cell cycle activity. Subsequently, cell cycle exit and entry into differentiation induces a shift from glycolysis to oxidative phosphorylation (OXPHOS), which becomes the primary energy-producing pathway in differentiated myotubes to generate increased ATP levels for their contractile activity. In a similar fashion, adult skeletal muscle consists of fiber types defined by their contractile properties and fiber-specific metabolic activity, and can transition between these cell states, remodeling their metabolism and gene expression patterns, to adapt to physiological demands (Bourdeau et al., 2018; Schiaffino and Reggiani, 2011). Lastly, a diseased cell state such as muscular dystrophy is associated with profound changes in metabolic activity, gene expression, and epigenetics indicative of their dysfunctional and degenerative condition (Fontes-Olivera et al., 2017; Groh et al., 2009; Hardee et al. 2021; Heydemann, 2018).

At the genome level, myoblasts and myotubes express cell state-specific gene programs that form their distinct identities and are established by histone modification patterns that depend on availability of intermediate metabolites (Nguyen et al., 2019; Nieborak and Schneider, 2018). Metabolites generated by glucose and lipid metabolism are required for histone acetylation in myogenic cells (Das et al., 2017; Pillon et al., 2022; Yucel et al., 2019). Changes in histone modifications in skeletal myofibers have also been documented during exercise, which has wide-ranging effects on metabolic reactions and the production of metabolites (Egan and Zierath, 2013, Hargreaves and Spriet, 2020; McGee et al., 2009). The molecular circuitry of this metabolo-epigenomic axis is best exemplified by the findings that metabolites such as acetyl-CoA and NAD+, whose levels fluctuate based on the prevailing metabolic activity in each myogenic cell state, regulate the activity of the histone acetyltransferase MYST1 and deacetylase SIRT1, respectively (Ryall et al., 2015; Sincennes et al., 2021). While these studies have revealed cooperation between metabolism and the epigenome, the mechanisms by which histone modifiers are coordinated with specific metabolic activities and cell state to remodel the chromatin landscape in myogenesis remains incomplete.

The imprinted *Dlk1-Dio3* ncRNA gene, encoding over 60 microRNAs (miRNAs), several long noncoding RNAs (lncRNAs), and a group of small nucleolar RNAs (snoRNAs), has emerged as a candidate regulator of metabolism and cell state. Expression levels of *Dlk1-Dio3* ncRNAs have been shown to correlate with changes in metabolic activity, including mitochondrial pathways, in the hematopoietic, pancreatic, and muscle lineages (Kameswaran et al., 2014; Labialle et al., 2014; Qian et al., 2016; Vu Hong et al., 2022; Wust et al., 2018). In addition, we previously reported that *Meg3*, a major lncRNA in this locus, regulates myoblast plasticity, but also noted mitochondrial defects in *Meg3*-deficient cells (Dill et al., 2021). Despite these intriguing observations, the studies are imperfect, as they have focused on individual or small groups of ncRNAs and have neglected to address the direct and collective role of the entire gene particularly since this cluster of distinct classes of ncRNAs is transcribed as a single, polycistronic primary RNA. Moreover, phenotypes have been attributed to the *Dlk1-Dio3* ncRNA cluster based on deletions in the intergenic differentially methylated region (IG-DMR) broadly affecting expression patterns across the one megabase (1 Mb) domain including the flanking *Dlk1* and *Dio3* and paternal *Rtl1* protein-coding genes, or deletion of the *Meg3* lncRNA coding region which also affects expression of numerous other ncRNAs in the cluster (Lin et al., 2003; Takahashi et al., 2009; Zhou et al., 2010; Zhu et al., 2019).

Investigating the function of the *Dlk1-Dio3* ncRNA cluster through targeted deletion of its entire gene body has been constrained by the massive size and complex organization of this >200 kilobase (kb) gene (Benetatos et al., 2013; da Rocha et al., 2008; Dill and Naya, 2018). As an alternate approach to determine the specific and collective role of this ncRNA cluster in the coordination of metabolism and cell state, we created two stable muscle lines harboring distinct deletions in the proximal promoter region upstream of *Meg3*, thereby restricting effects on expression to the ncRNA cluster. Indeed, changes in *Dlk1-Dio3* ncRNA expression were highly specific, resulting in either massive upregulation or downregulation of the entire cluster but not neighboring genes in the domain. Both *Dlk1-Dio3* ncRNA promoter mutant lines, with contrasting levels of expression, displayed impaired myotube formation and OXPHOS activity. Furthermore, these defects correlated with genome-wide changes in chromatin accessibility and histone methylation patterns that dramatically altered their cell state, prematurely priming proliferating myoblasts for differentiation.

## RESULTS

### Deletions in the *Meg3* proximal promoter region in myoblasts cause differential effects on *Dlk1-Dio3* noncoding RNA expression

Defining the regulatory function of the entire cluster of ncRNAs expressed from the imprinted *Dlk1-Dio3* locus has been hindered by the technical challenges of deleting this >200 kb chromosomal region (Benetatos et al., 2013; da Rocha et al., 2008; Dill and Naya, 2018). While several groups have removed genomic sequences corresponding to small clusters of ncRNAs within the *Dlk1-Dio3* gene body, these approaches have failed to directly address the collective function of the ncRNA cluster (Gao et al., 2015; Labialle et al., 2014). This is not an insignificant consideration as the gene structure of this massive ncRNA cluster, encoding a diversity of ncRNAs, has been conserved in mammals and is transcriptionally regulated by a proximal promoter immediately upstream of *Meg3*, suggesting their coexpression is biologically important (Edwards et al., 2008; Kircher et al., 2008; Snyder et al., 2013; Zhao et al., 2005).

To overcome this obstacle CRISPR/Cas9 was used to create a deletion in the *Meg3* proximal promoter in C2C12 myoblasts that would be expected to disrupt transcription and result in downregulated expression of the entire ncRNA cluster. Guide RNAs (gRNAs) were designed to delete ∼300 base pairs upstream from the TATA sequence (Figure 1A). C2C12 myoblasts were co-transfected with either Cas9 alone or Cas9 and promoter-specific guide RNAs followed by GFP sorting to isolate transfected cells. From a serially diluted pool of GFP-positive myoblasts, single cells were seeded onto 96-well plates and grown without antibiotic selection (to prevent integration of Cas9/gRNA plasmids) followed by extraction of genomic DNA for PCR analysis of the *Meg3* promoter region. As shown in Figure 1B (left panel), one clone designated Meg3-CAS (proximal *<u>c</u>is*-<u>a</u>cting <u>s</u>equence) out of hundreds screened, displayed a single, smaller DNA band compared to control myoblasts, pX (Cas9 plasmid transfection only). Sequencing performed on this DNA fragment confirmed a deletion between nucleotides −339 and −43 (Fig. 1B, right panel). Notably, the TATA box and overlapping MEF2 site (Snyder et al., 2013) located near position −37 were still present, likely reflecting an imperfect cleavage by Cas9 at the location of the 3’ gRNA primer.

**Fig. 1.**
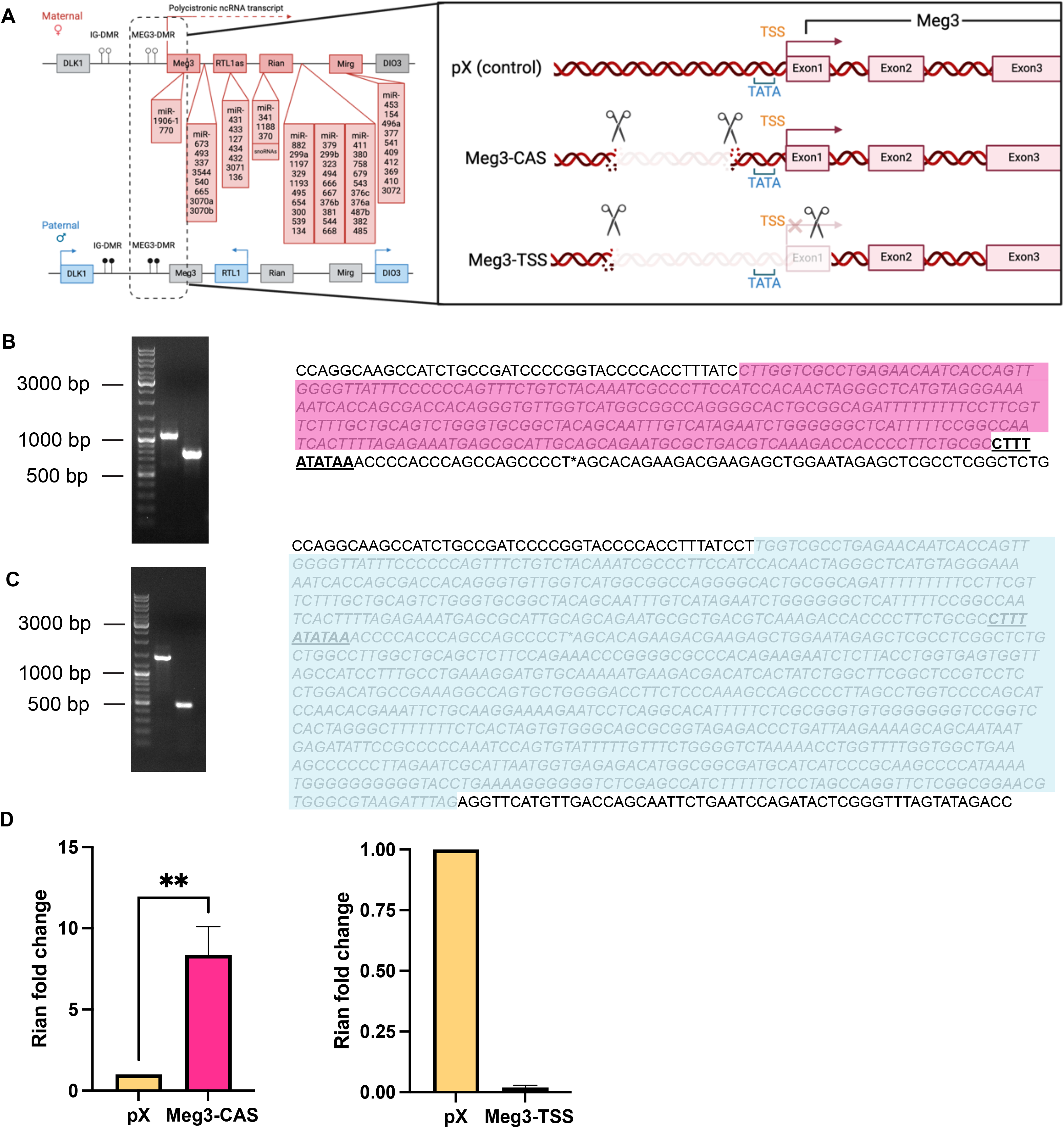
CRISPR-Cas9 induced deletions in the *Meg3* proximal regulatory region in C2C12 myoblasts. (A) Schematic of regions deleted in C2C12 cells using CRISPR-Cas9 gene editing. Guide RNAs were designed to target two regions in the *Meg3* promoter region resulting in two stable clones, Meg3-CAS and Meg3-TSS. (B) Confirmation of deletion in Meg3-CAS cells present in Meg3 proximal regulatory region. (Left) Gel electrophoresis of genomic DNA amplicons (from left to right): DNA ladder, PCR amplicon from C2C12 transfected with pX458 vector (pX cells), PCR amplicon from C2C12 transfected with pairs of pX458/guide RNA constructs (Meg3-CAS cells). pX cells display the 1.1 kb band of the wild-type Meg3 proximal promoter. Meg-CAS C2C12 cells display an approximately ∼800 bp band, indicating a ∼300 bp homozygous deletion in the Meg3 proximal regulatory region which was confirmed by sequencing (right; grey italics and pink highlighted). Note the intact TATA box (bold and underlined). (C) Confirmation of deletion in Meg3-TSS cells present in Meg3 proximal regulatory region. (Left) Gel of C2C12 genotyping amplicons (from left to right): DNA Ladder, genotyping PCR product from C2C12 transfected with pX458 vector (pX cells), genotyping PCR product from C2C12 transfected with pairs of pX458/other guide RNA constructs (Meg3-TSS cells). pX cells display the 1.469 kb band predicted for amplification of the wild-type Meg3 proximal promoter allele. Meg-TSS C2C12 cells display an approximately 519 bp band, indicating an approximately 950 bp homozygous deletion in the Meg3 proximal regulatory region including the TSS and TATA box, shown as the gray italics and blue highlighted sequencing (right). (D) qPCR quantification of *Dlk1-Dio3* ncRNA Rian confirms overexpression of *Dlk1-Dio3* ncRNA in Meg3-CAS cell and reduction of *Dlk1-Dio3* ncRNA in Meg3-TSS compared to pX control cells (n=3).

To determine the effect this deletion had on *Dlk1-Dio3* ncRNA expression in Meg3-CAS cells we examined transcript levels of *Rian*, a representative lncRNA located in the middle of the polycistronic primary RNA transcript, which is readily detected in C2C12 myoblasts. Unexpectedly, *Rian* expression was significantly increased in Meg3-CAS myoblasts relative to pX control cells (Figure 1D, left bar graph). These results suggest that a repressive *cis*-acting sequence(s) in the proximal promoter was deleted resulting in greatly enhanced transcription.

To create another myoblast line with downregulated expression of all *Dlk1-Dio3* ncRNAs a subsequent CRISPR/Cas9-mediated screen was performed as described above using a different gRNA primer pair designed to create a larger deletion encompassing the proximal promoter sequence, TATA box, transcription start site (TSS), and a short stretch of non-functional, poorly conserved nucleotides at the 5’-end of murine *Meg3* (−339 to +585) (Fig. 1A) (Uroda et al., 2019). This screen identified a single clone, designated Meg3-TSS, harboring a ∼900 bp homozygous deletion (Fig. 1C, left and right panels). In this instance, expression of *Rian* in Meg3-TSS myoblasts was substantially reduced relative to pX control cells (Fig. 1D, right bar graph). The creation of these novel mutant myoblast lines harboring two distinct deletions in the *Meg3* promoter region - resulting in super-physiological and downregulated expression of *Dlk1-Dio3* ncRNAs - enabled us to precisely and specifically define the function of the entire ncRNA cluster and how changes in its dosage impact muscle cell state and differentiation.

### Meg3-CAS and Meg3-TSS myoblasts with altered *Dlk1-Dio3* ncRNA dosage display impaired cell state and differentiation

Genomic imprinting of the *Dlk1-Dio3* ncRNA locus suggests that dosage is critical for cell function (Weinberg-Shukron et al., 2023). Consistent with this notion, we noted that both Meg3-CAS and Meg3-TSS proliferating myoblasts displayed considerably slower growth than pX controls, necessitating 12-24 hrs longer to reach the same density (data not shown). To determine how changes in *Dlk1-Dio3* ncRNA dosage affect muscle differentiation, Meg3-CAS, Meg3-TSS, and pX control myoblasts were induced to differentiate. Between 24 and 48 hrs after inducing differentiation, there was decreased confluency in Meg3-CAS cells, as well as an increase in detached cells. By day 3 of differentiation, most Meg3-CAS myoblasts formed smaller, less elongated (stubbier) myotubes compared to pX myotubes (Fig. 2A, middle panels). Meg3-CAS myotubes maintained their growth and viability through day 7 of differentiation but still appeared smaller compared to pX controls (data not shown). By contrast, Meg3-TSS myoblasts formed thinner and more elongated myotubes than Meg3-CAS mutants, giving the appearance of a web-like network of multinucleated myotubes, yet their morphology was clearly abnormal compared to pX myotubes (Fig. 2A, right panels). These results indicate that myoblasts are exquisitely sensitive to dosage of *Dlk1-Dio3* ncRNAs and that their dysregulated expression (up or down) impairs proper myotube formation.

**Fig. 2.**
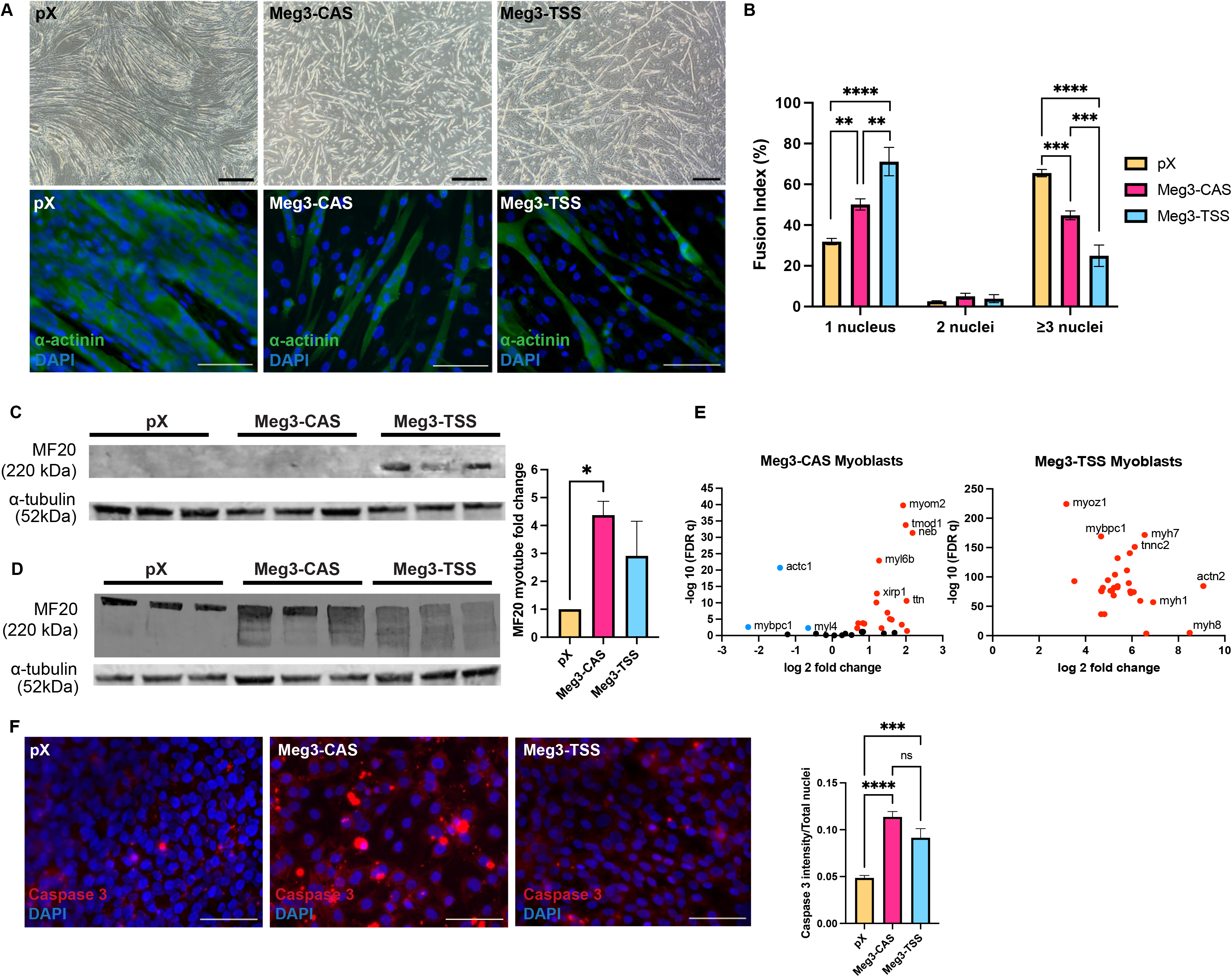
Altered *Dlk1-Dio3* ncRNA expression severely impairs myotube formation in Meg3-CAS and Meg3-TSS mutant cells. (A) (Top panel) Brightfield images on day 3 of differentiation (D3). The pX control cells display long connected myotube formation whereas Meg3-CAS and Meg3-TSS myotubes are shorter with altered morphologies. (Bottom panel) Immunocytochemistry images on D3 were used to quantity fusion index. Myotubes were stained using α-actinin (green) to define cell area and DAPI (blue) for nuclei. Scale bar = 500 μm. (B) The fusion index was calculated based on the percentage of nuclei within cells, delineated by α-actinin expression, containing one nucleus, two nuclei, and more than three nuclei, with each category representing a proportion of the total nuclei within the cell border totaling 100%. Fusion into mature myotubes (≥3 nuclei) was reduced in both mutant cell lines compared to pX myotubes (n=4). Scale bar = 100 μm. (C) Western blot quantification of MF20 (normalized to α-tubulin) for myoblasts demonstrates premature expression of myosin heavy chain in Meg3-TSS cells (n=3). (D) Western blot quantification of MF20 (normalized to α-tubulin) for myotubes at D3 reveals increased expression of MF20 in Meg3-CAS but not Meg3-TSS myotubes compared to pX myotubes. E) (Left) Volcano plot depicting log 2 fold change of sarcomeric genes from RNA-seq in Meg3-CAS myoblasts compared to pX controls where red dots represent upregulated genes with a FDRq <0.05 and blue dots represent downregulated genes with a a FDRq <0.05. (Right) Volcano plot depicting log 2 fold change of sarcomeric genes from RNA-seq in Meg3-TSS myoblasts compared to pX controls where red dots represent upregulated genes with a FDRq <0.05. (F) Immunocytochemistry images for Caspase 3 one day after changing to differentiation media (n=4). Scale bar = 100 μm. Quantification of Caspase 3 intensity (red) per field of view normalized to total number of nuclei as shown by DAPI in blue demonstrates increased Caspase 3 in both mutant cell lines compared to pX control cells upon differentiation (n=3).

Meg3-CAS and Meg3-TSS mutants were subsequently evaluated for fusion index, an indicator of robust formation of multinucleated myotubes. On differentiation day 3 both Meg3-CAS and Meg3-TSS mutants had a greater percentage of α-actinin positive myotubes with one nucleus compared to pX control myotubes (Fig. 2B). Conversely, both mutants had a significantly smaller proportion of myotubes with 3 or more nuclei, indicative of impaired myoblast fusion (Fig. 2B). Notably, whereas the Meg3-CAS mutant line had a similar distribution of myotubes with one nucleus or greater than 3 nuclei, Meg3-TSS mutant myotubes had a higher percentage of mononucleated cells relative to those with 3 or more nuclei (Fig. 2B).

To corroborate the myotube formation defects, expression of sarcomeric myosin heavy chain 4 (MYH4), a sensitive indicator of differentiation, was examined. Paradoxically, despite the abnormalities in myotube formation, both Meg3-CAS and Meg3-TSS myotubes on day 3 of differentiation expressed substantially higher levels of MYH4 protein compared to pX myotubes (Fig. 2D and bar graph). This prompted us to consider whether MYH4 was aberrantly expressed in proliferating myoblasts which do not express this myosin isoform. As shown in Fig. 2C, Meg3-TSS but not Meg3-CAS or pX proliferating myoblasts expressed MYH4 protein. The abnormal expression of MYH4 in mutant myoblasts raised the possibility that changes in *Dlk1-Dio3* ncRNA dosage altered myoblast cell state such that genes associated with differentiation are inappropriately expressed in proliferating cells. To investigate this notion, we examined RNA-sequencing (RNA-seq) data for additional upregulated transcripts encoding sarcomere and other muscle structural proteins in Meg3-CAS and Meg3-TSS mutants. As shown in Fig. 2E, dozens of sarcomeric and other structural transcripts were significantly upregulated in both mutant myoblasts. Interestingly, RNA-seq also revealed that expression of myogenic transcriptional regulators *myogenin* and *Mef2C*, but not *MyoD* were upregulated in both Meg3-CAS and Meg3-TSS myoblasts (Supplemental File). Taken together, induction of MYH4 and numerous other sarcomeric and structural transcripts, and myogenic regulatory factors in Meg3-CAS and Meg3-TSS myoblasts strongly suggests that myoblasts are exquisitely sensitive to dosage of *Dlk1-Dio3* ncRNAs and deviation from these normal levels, an increase or decrease, provokes remodeling of myoblasts to a differentiated-like cell state.

As noted above, differentiation experiments also revealed numerous detached cells in the early stages of this process (12-24 hours after inducing differentiation) in Meg3-CAS and Meg3-TSS myoblasts indicating cell viability was affected by the dysregulated expression of *Dlk1-Dio3* ncRNAs. Therefore, we examined programmed cell death by measuring caspase-3 expression at this time point. As shown in Fig. 2F, both mutant lines showed significantly higher caspase-3 immunoreactivity relative to pX control cells 12 hours post-induction. Thus, changes in dosage of *Dlk1-Dio3* ncRNAs in myoblasts induced to differentiate caused increased cell death which could explain, in part, the inadequate formation of multinucleated myotubes.

### Meg3-CAS and Meg3-TSS mutant cell lines display largely distinct but overlapping dysregulated cellular processes

To delve deeper into the molecular mechanisms that lead to abnormal myotube formation in Meg3-CAS and Meg3-TSS cells, we performed RNA-seq in both mutant myoblasts and myotubes. A broad comparison of the total number of dysregulated transcripts (FDR *q* < 0.05, > 2-fold change) in Meg3-CAS and Meg3-TSS mutants revealed the extent to which myoblasts and myotubes differentially responded to changes in *Dlk1-Dio3* ncRNA dosage (Figure 3A). Overexpression of the *Dlk1-Dio3* ncRNA cluster (Meg3-CAS) triggered significantly greater downregulation of gene expression (1334 genes) than upregulation (980 genes) in proliferating myoblasts compared to differentiated myotubes, which had similar numbers of up-and down-regulated transcripts (2120 down, 2117 up; Fisher’s exact *p* = 3.5 x 10^-9^). Conversely, reduced expression of *Dlk1-Dio3* ncRNAs (Meg3-TSS) was associated with significantly more downregulation (2950 genes) than upregulation (2552 genes) in myotubes compared to myoblasts (1536 down, 1644 up; Fisher’s exact *p* = 1.8 x 10^-6^).

**Fig. 3.**
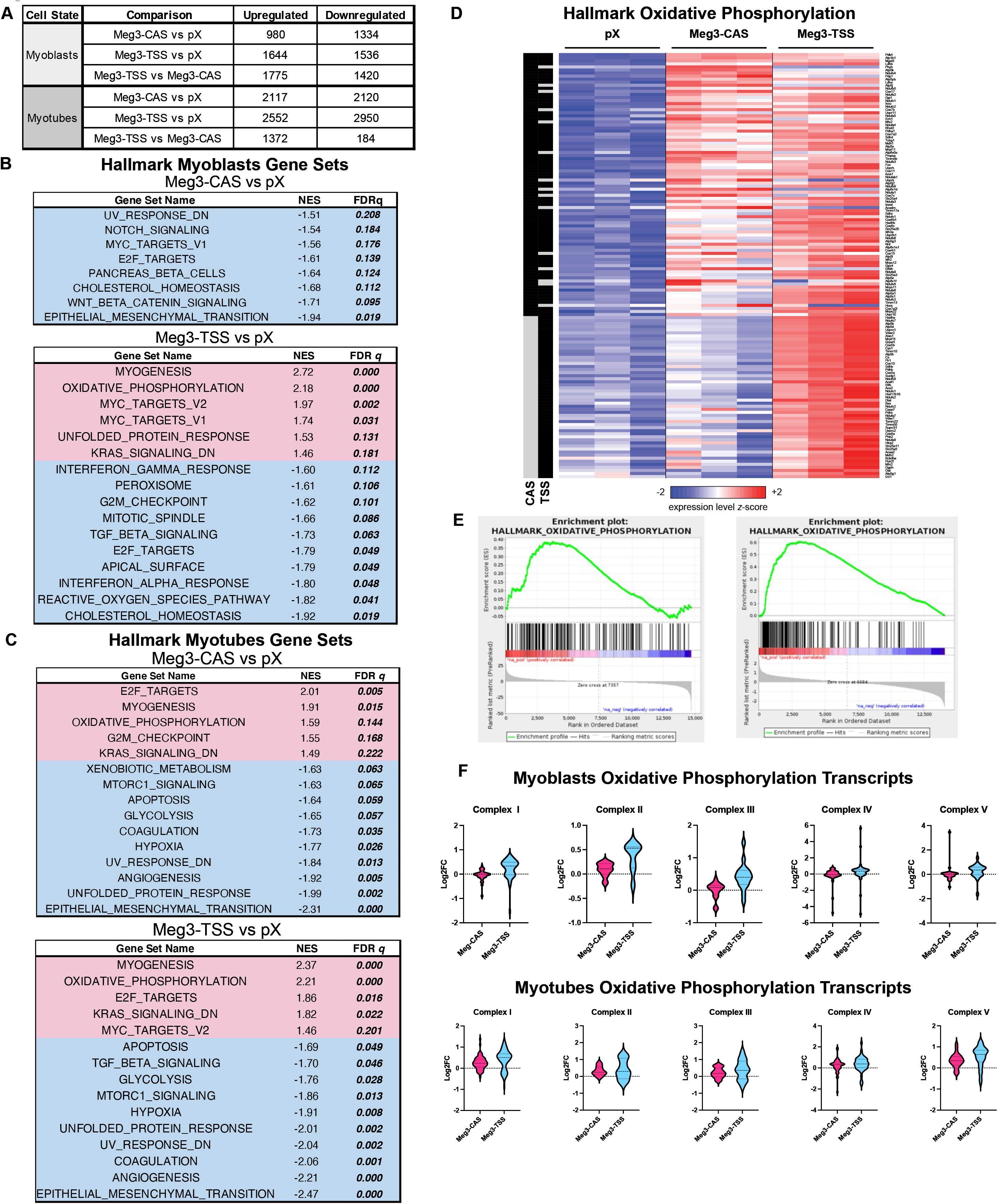
Transcripts belonging to oxidative phosphorylation are significantly enriched in Meg3-CAS and Meg3-TSS mutant myoblasts and myotubes. (A) Summary table of RNA-seq dysregulated transcripts with FDR q < 0.05 and fold change > 2 (upregulated) or < −2 (downregulated) demonstrates differential gene expression patterns in mutant cell lines. (B-C) Hallmark gene set enrichment analysis genes for the top ten up and down-regulated Hallmark gene sets that meet the FDRq <0.25 threshold for myoblasts (B) and myotubes (C). (D) Heatmap showing differential gene expression for the Hallmark Oxidative Phosphorylation gene set in which Meg3-CAS and Meg3-TSS myotubes are upregulated compared to pX myotubes. (E) Enrichment plots showing upregulation in the Hallmark Oxidative Phosphorylation gene set for both Meg3-CAS and Meg3-TSS myotubes. (F) Violin plots of RNA-seq data for MitoCarta3.0 gene sets for each complex involved in oxidative phosphorylation generally show a positive fold change trend for both Meg3-CAS and Meg3-TSS myoblasts which becomes more pronounced at the myotube stage compared to pX control cells.

We then used Gene Set Enrichment Analysis (GSEA) with the MSigDB Hallmark collection of gene sets to identify pathways that were coordinately up- or down-regulated between each mutant and pX control within myoblasts (Supplementary File 1). There were no gene sets that were coordinately upregulated in Meg3-CAS myoblasts at a stringent false discovery rate (FDR *q* < 0.05), although several pathways were significant at the more lenient threshold of FDR *q* < 0.25, including KRAS signaling, TNFα/NF-κB signaling, and estrogen response (Fig. 3B). In contrast, many gene sets were significantly coordinately downregulated (FDR *q* < 0.05) in Meg3-CAS myoblasts, including epithelial-mesenchymal transition (EMT), Wnt/β-catenin signaling, cholesterol biosynthesis, E2F targets, and Notch signaling (Fig. 3B). In the Meg3-TSS myoblasts, pathways with significant coordinate upregulation included myogenesis, OXPHOS, MYC targets, and the unfolded protein response (UPR), and those with significant coordinate downregulation pertained to cholesterol biosynthesis, reactive oxygen species (ROS), E2F targets, TGFβ signaling, and EMT. Notably, the Meg3-CAS and Meg3-TSS myoblasts shared a few downregulated processes in common, including cholesterol homeostasis, E2F targets, and EMT.

The same GSEA analysis was then applied to myotubes, which showed that there was strong overlap between the Meg3-CAS and Meg3-TSS cell lines. In both mutants, myogenesis, OXPHOS, and E2F targets were significantly coordinately upregulated compared to pX control, whereas EMT, UPR, hypoxia, and glycolysis were significantly coordinately downregulated (FDR *q* < 0.05, Fig. 3C). G2/M checkpoint genes were also strongly upregulated (FDR *q* < 0.1) in both cell lines, although some pathways were mutation-specific: MYC targets were upregulated and TGF-β signaling was downregulated in the Meg3-TSS, but not the Meg3-CAS, cell line (Fig. 3C). Taken together, these results suggest that a dramatic remodeling of metabolism and cell state occurs in muscle cells with altered dosage of *Dlk1-Dio3* ncRNAs.

### Alterations in oxidative phosphorylation in Meg3-CAS and Meg3-TSS mutant cells

Based on the pathway analyses above we explored the possibility that OXPHOS is a key target of *Dlk1-Dio3* ncRNAs in both mutant lines because it emerged as one of the most strongly dysregulated processes across the two genotypes in myotubes (Fig. 3D and E). Moreover, it provided us an opportunity to gain insight into the regulatory network coordinating metabolism, cell state, and muscle differentiation.

Further detailed bioinformatic analysis revealed consistent upregulation of the expression of numerous nuclear- and mitochondrial-encoded transcripts encoding components of the electron transport chain (ETC) (Fig. 3F). To determine whether ETC protein expression was similarly affected, we analyzed the levels of selected subunits from each ETC complex in enriched mitochondrial fractions from both mutant lines (Complex I [NADH dehydrogenase (ubiquinone) 1 beta subcomplex subunit 8 (NDUFB8)], II [succinate dehydrogenase B (SDHB)], III [mitochondrial cytochrome c oxidase 1 (MT-CO1)], IV [ubiquinol-cytochrome C reductase rore protein 2 (UQCRC2)], and V [ATP synthase 5A (ATP5A)]) (Fig. 4A). In myoblasts, we observed a significant decrease in nuclear-encoded SDHB in Meg3-TSS myoblasts compared to pX control and Meg3-CAS cells (Fig. 4A, left bar graph). In contrast, protein levels of a mitochondrial-encoded subunit of cytochrome oxidase Complex IV, MT-CO1, were significantly upregulated in Meg3-CAS but not Meg3-TSS myoblasts (Fig. 4A, right bar graph). Examination of OXPHOS proteins in myotubes revealed a dramatically different profile in *Dlk1-Dio3* ncRNA mutant cells. Unlike myoblasts, both Meg3-CAS and Meg3-TSS myotubes had significantly upregulated SDHB expression (Fig. 4B, left bar graph). MT-CO1 continued to be significantly upregulated in Meg3-CAS but not Meg3-TSS myotubes (Fig. 4B, right bar graph). Expression of the remaining OXPHOS subunits was not significantly different in either mutant myoblasts or myotubes compared to pX controls (Supplemental Figure S1).

**Fig. 4.**
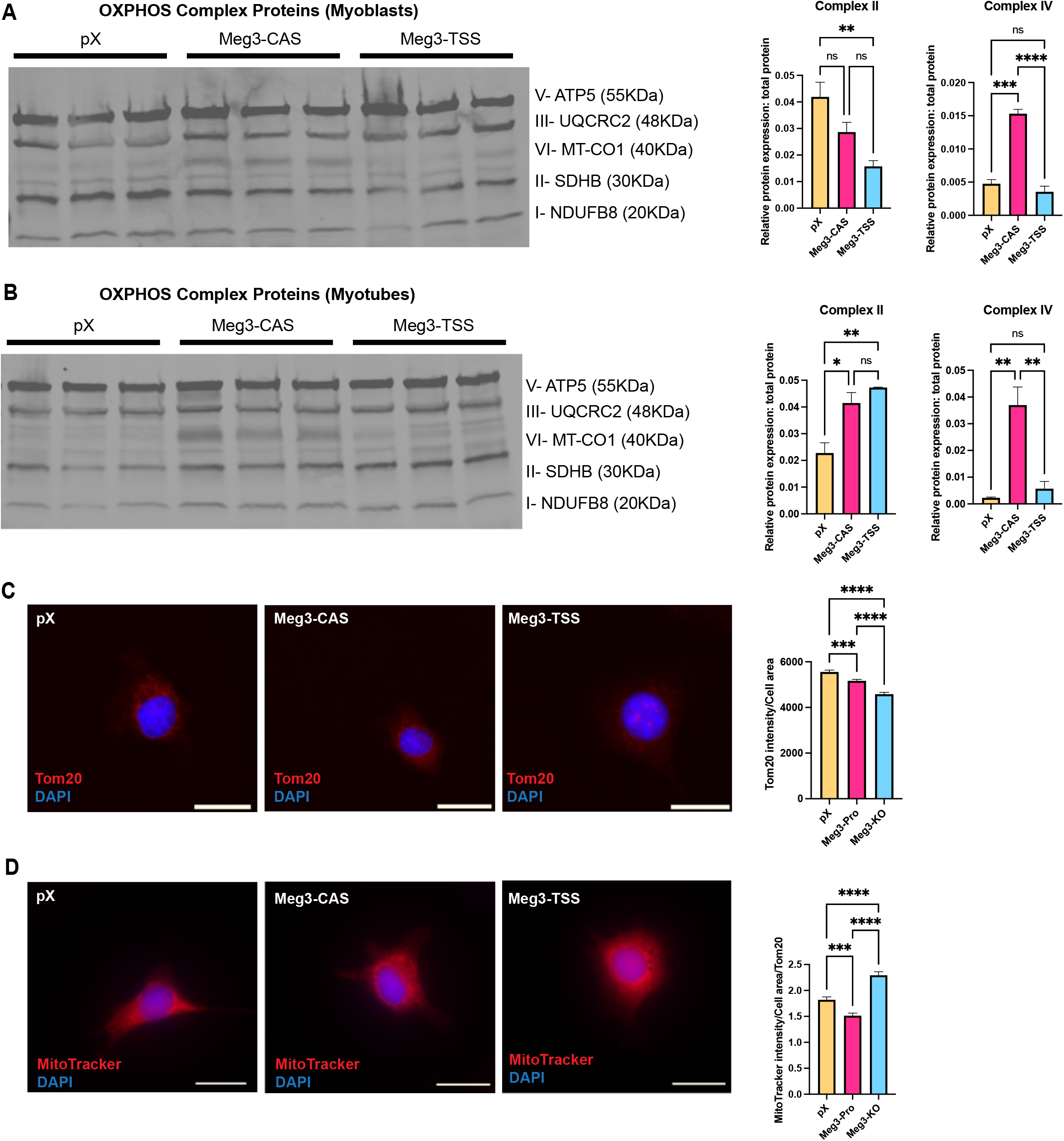
Altered dosage of *Dlk1-Dio3* ncRNAs causes changes to mitochondrial biomass and membrane potential in cells with altered *Dlk1-Dio3* ncRNA expression. (A) Western blot quantification of select OXPHOS complex subunits (normalized to total protein) for myoblasts demonstrates lower Complex II (SDHB) expression in Meg3-TSS cells compared to pX control cells and increased expression of Complex IV (MT-CO1) in Meg3-CAS cells compared to both pX control and Meg3-TSS cells (n=3). (B) Western blot quantification of select OXPHOS complex subunits (normalized to total protein) for myotubes demonstrates increased Complex II (SDHB) expression in both mutant lines compared to pX control cells and increased expression of Complex IV (MT-CO1) in Meg3-CAS cells compared to both pX control and Meg3-TSS cells (n=3). (C) Immunocytochemistry for Tom 20 was used to assess mitochondrial mass in myoblasts in which Tom20 intensity was normalized to cell area defined α-actinin (green channel not shown). Both mutant cell lines had lower mitochondrial mass compared to the control (n=4). Scale bar = 25 μm. (D) Myoblasts were incubated with MitoTracker Red CMXRos for 30 minutes, as a method to indirectly assess mitochondrial membrane potential, and co-stained with α-actinin. Quantification of MitoTracker signal was confined to α-actinin + cells and normalized to average mitochondrial mass per cell line (n=4). MitoTracker quantification revealed a reduction in mitochondrial membrane potential for Meg3-CAS cells compared to both pX and Meg3-TSS, while pX cells had reduced membrane potential compared to Meg3-TSS cells (n=4). Scale bar = 25 μm.

We next sought to investigate mitochondrial mass and membrane potential in *Dlk1-Dio3* ncRNA mutant myoblasts. Initially, mitochondrial mass was determined by quantifying the immunofluorescence signal of the mitochondrial outer membrane receptor, Tom20, within a myoblast as defined by the α-actinin positive cell area (Fig. 4C, upper panels). Both Meg3-CAS and Meg3-TSS had a modest but significant reduction in Tom20 expression indicating decreased mitochondrial mass in both mutants (Fig. 4C, upper right bar graph). Subsequently, membrane potential was assessed through examination of MitoTracker CMXRos. As shown in Fig. 4D (lower panels), MitoTracker Red CMXRos signal intensity was significantly reduced in Meg3-CAS but significantly increased in Meg3-TSS myoblasts reflecting diminished and augmented mitochondrial membrane potential, respectively, relative to pX controls (Fig. 4D, lower right bar graph).

Given the dysregulated expression of OXPHOS subunits and altered mitochondrial membrane potential, mitochondrial respirometry was performed on Meg3-CAS and Meg3-TSS myoblasts. Proliferating Meg3-CAS but not Meg3-TSS myoblasts displayed a significant increase in basal oxygen consumption rate (OCR), i.e. respiration (Fig. 5A and B). The most dramatic increase in OCR measured in Meg3-CAS myoblasts occurred after administering the protonophore FCCP, a mitochondrial uncoupler (Fig. 5A and C). Further, following Complex I inhibition, ATP-linked respiration was calculated and observed to be highest in Meg3-CAS compared to Meg3-TSS and pX myoblasts (Fig. 5A and D). These results are consistent with the upregulated expression of the Complex IV subunit MT-CO1 in Meg3-CAS but not Meg3-TSS mutants (Fig. 4A and B). Additional parameters were calculated based on the respirometry measurements to better understand respiration in mutant myoblasts. Proton leak was found to be significantly increased in Meg3-CAS myoblasts but decreased in Meg3-TSS mutants (Fig. 5E). The increased proton leak in Meg3-CAS mutants was surprising given their increased respiration. Spare respiratory capacity, reflecting the extent of maximal respiration over basal levels, was also significantly higher in Meg3-CAS myoblasts but lower in Meg3-TSS (Fig. 5F). Similarly, coupling efficiency (ATP production rate/basal respiration) was significantly higher in Meg3-CAS cells compared to both Meg3-TSS and pX controls (Fig. 5G). Non-mitochondrial OCR was also significantly higher in Meg3-CAS, but significantly lower in Meg3-TSS mutant myoblasts (Supplemental Figure S2). For all respiration measurements, Meg3-TSS myoblasts had significantly decreased OCR compared to pX controls. These results clearly indicate that mitochondrial respiration is exquisitely sensitive and differentially affected by dosage of *Dlk1-Dio3* ncRNAs such that respiration is enhanced by their overexpression but dampened by their reduced expression.

**Fig. 5.**
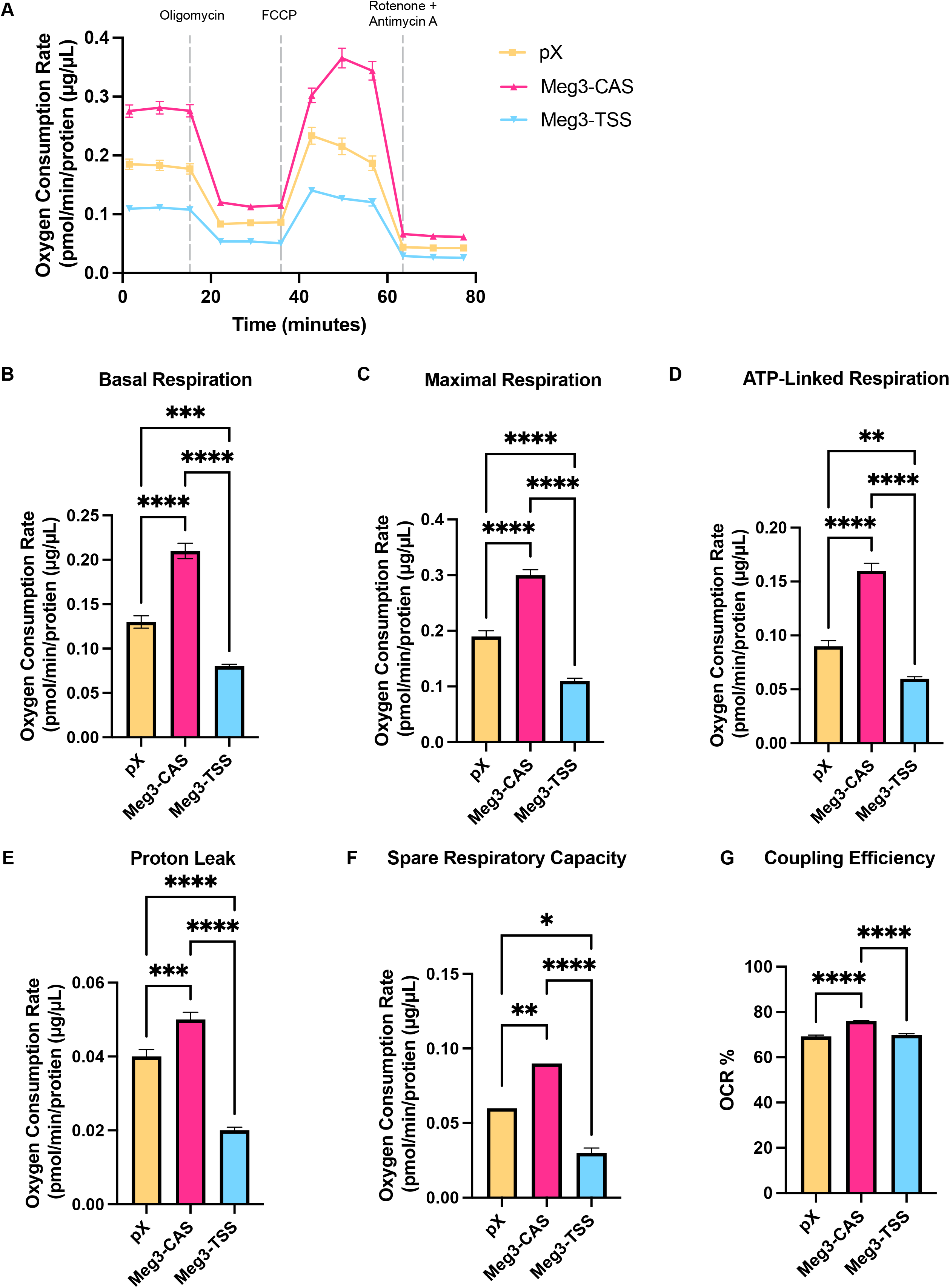
Altered mitochondrial respiration in *Dlk1-Dio3* ncRNA mutant cells. (A) OCR of pX, Meg3-CAS, and Meg3-TSS cells in response to 1.5 μM oligomycin, 1 μM FCCP, 0.5 μM Rotenone and Antimycin A. (B) Basal respiration was quantified prior to injection of any inhibitors. (C) Maximal respiration was determined following the addition of FCCP. (D) ATP-linked respiration was quantified as the difference between basal respiration and proton leak. (E) Proton leak was determined following the addition of oligomycin. (F) Spare respiratory capacity was calculated as the difference between maximal and basal respiration. (G) Coupling efficiency was calculated as ATP-production rate divided by basal respiration.

### Genome-wide changes in chromatin accessibility and histone methylation patterns in Meg3-CAS and Meg3-TSS mutant cells

Having established defects in mitochondrial metabolism in *Dlk1-Dio3* ncRNA mutant cells and given the essential requirement of intermediate metabolites for histone modifications, we reasoned this would cause global alterations in chromatin structure.

To assess genome-wide changes in chromatin accessibility we subjected chromatin from Meg3-CAS and Meg3-TSS myoblasts and myotubes to ATAC-sequencing (ATAC-seq). Principal component analysis (PCA) of the ATAC-seq data revealed genotype-specific clustering across the three groups (Fig. 6A). Curiously, unlike the readily separable clustering between myoblasts and myotubes in the pX and Meg3-TSS groups, the tight clustering of Meg3-CAS myoblasts and myotubes suggested these distinct cell states have similar chromatin accessibility. When comparing the overall number of shared and unique ATAC-seq peaks, Meg3-CAS and Meg3-TSS myoblasts and myotubes were substantially different from each other and pX controls, consistent with their distinct phenotypes (Fig. 6B). Based on these observations we examined chromatin accessibility profiles in both mutant lines in greater detail focusing on OXPHOS as this was the most significantly enriched pathway. As shown by the volcano plots in Fig. 6C-F, pX control myoblast and myotube genomes including OXPHOS loci (black dots) displayed a greater degree of chromatin accessibility than both Meg3-CAS and Meg3-TSS mutants, though this difference was less pronounced in Meg3-TSS mutants. This was an unexpected result given the upregulated expression of OXPHOS transcripts in both mutant lines but could relate to the regulatory complexity at these loci (see Discussion). In addition to OXPHOS we also examined EMT in our ATAC-seq data set given this was the most significantly downregulated pathway in both mutant lines. In this instance, both *Dlk1-Dio3* ncRNA mutant lines displayed reduced chromatin accessibility demonstrating concordance between the transcriptome and chromatin accessibility (Supplemental Figure S3).

**Fig. 6.**
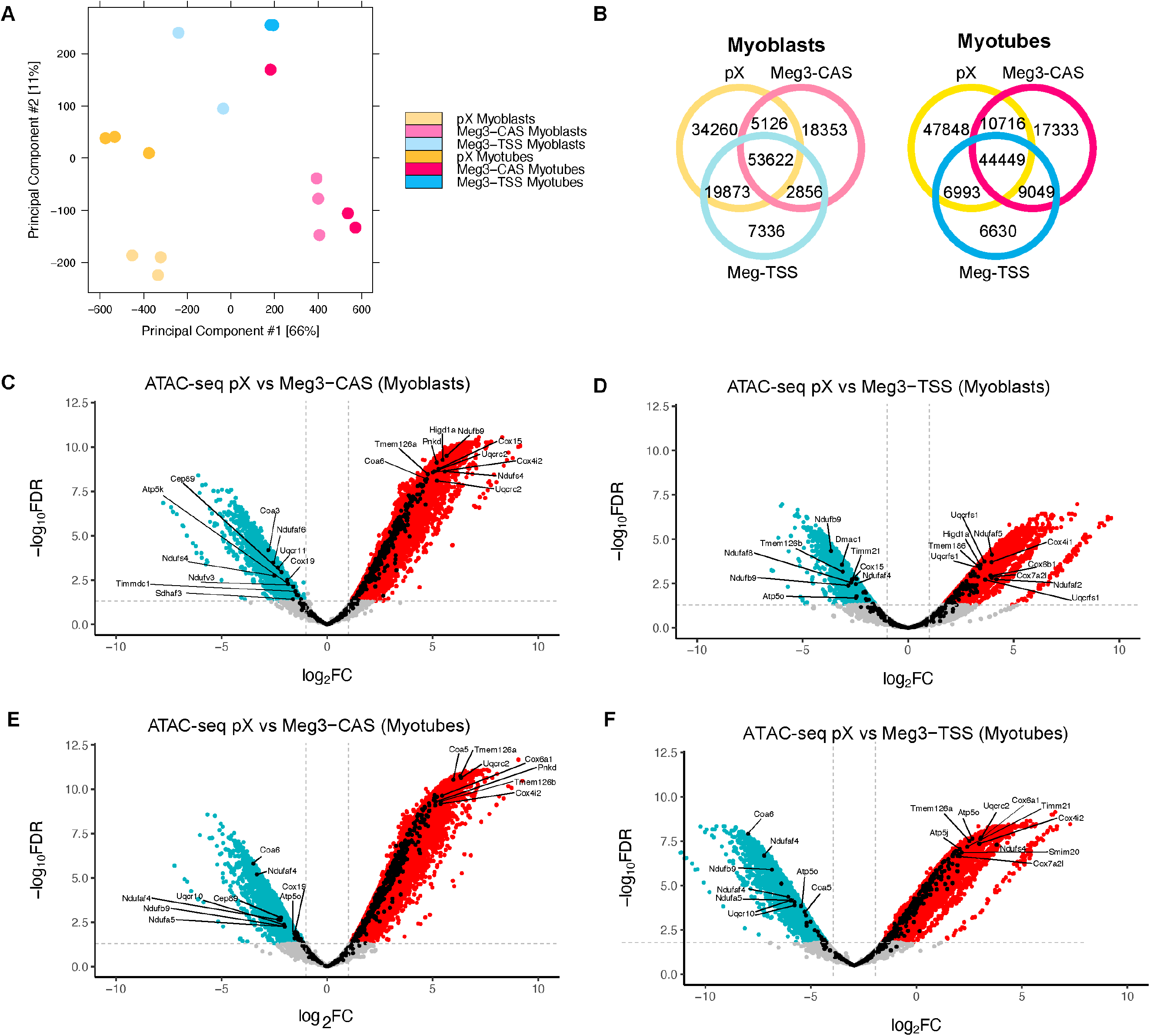
Differential chromatin accessibility modulated by the *Dlk1-Dio3* ncRNA cluster. (A) Principal component analysis from ATAC-seq data reveals distinct differences between cell lines. (B) Venn diagram of ATAC-seq peaks for each cell line pX, Meg3-CAS and Meg3-TSS and number of peaks in common for myoblasts and myotubes. (C) Volcano plot of ATAC-seq data comparing differentially open regions of chromatin in pX compared to Meg3-CAS myoblasts, with OXPHOS genes denoted in black (Top 10 unique OXPHOS genes labeled). (D) Volcano plot of ATAC-seq data comparing differentially open regions of chromatin in pX compared to Meg3-TSS myoblasts, with OXPHOS genes denoted in black (Top 10 unique OXPHOS genes labeled). (E) Volcano plot of ATAC-seq data comparing differentially open regions of chromatin in pX compared to Meg3-CAS myotubes, with OXPHOS genes denoted in black (Top 10 unique OXPHOS genes labeled). (F) Volcano plot of ATAC-seq data comparing differentially open regions of chromatin in pX compared to Meg3-TSS myotubes, with OXPHOS genes denoted in black (Top 10 unique OXPHOS genes labeled).

We next examined histone methylation in both mutant cell lines by chromatin immunoprecipitation using CUT&RUN sequencing. Initially, we focused on the repressive histone modification H3K27me3 given the established interaction between PRC2 and *Meg3* and other locus lncRNAs, *Rian* and *Mirg* (Kaneko et al., 2014; Zhao et al., 2010). As shown in Fig. 7A, the global H3K27me3 pattern was largely indistinguishable between the two mutant myoblast genomes, but it was greatly increased in Meg3-CAS myotubes compared to pX controls and Meg3-TSS myotubes, likely reflecting enhanced recruitment of PRC2 at target loci due to the massive upregulation of *Dlk1-Dio3* lncRNAs in this mutant background.

**Fig. 7.**
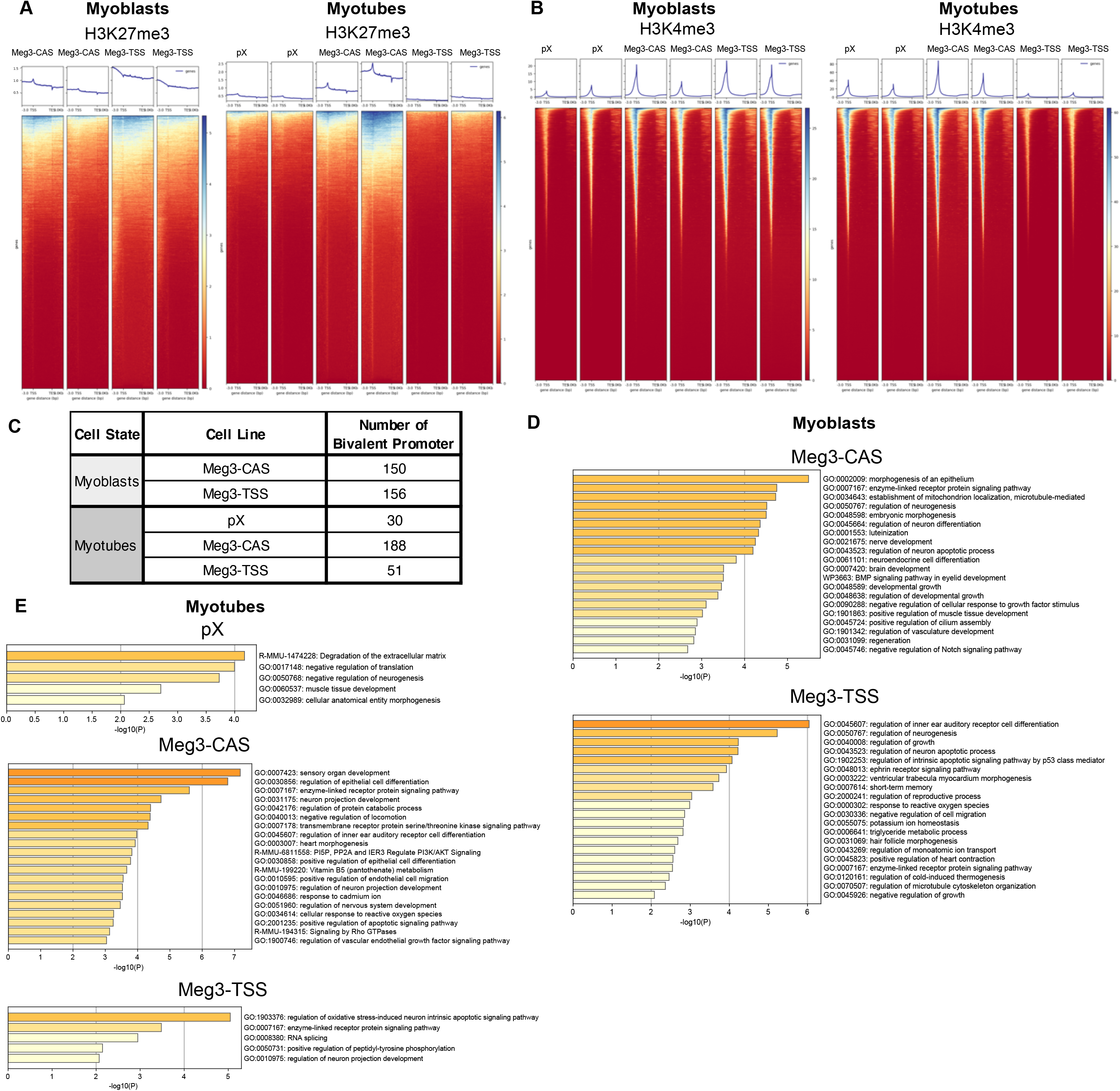
Distinct histone methylation patterns are dependent on *Dlk1-Dio3* ncRNA dosage. (A) CUT&RUN data from H3K27me3 showing no difference in genome-wide histone modification between Meg3-CAS and Meg3-TSS myoblasts (left heatmaps). The H3K27me3 profile was increased in Meg3-CAS myotubes compared to pX and Meg3-TSS myotubes. (B) CUT&RUN data from H3K4me3 for myoblasts showed increased H3K4me3 in mutant cell lines compared to pX control (left heatmaps). Whereas, the H3K4me3 profile for myotubes was increased for Meg3-CAS compared to pX control and decreased in Meg3-TSS (right heatmaps). (C) Summary table of bivalent promoter analysis for myoblasts and myotubes. (D) GO enrichment analysis for bivalent promoters in mutant myoblasts. (D) GO enrichment analysis for bivalent promoters in pX, Meg3-CAS, and Meg3-TSS myotubes.

Having profiled a repressive chromatin mark we were also interested in evaluating H3K4me3, a histone mark associated with transcriptional activation, to obtain a fuller picture of *Dlk1-Dio3* ncRNA dosage effects on epigenetic regulation in muscle cells. H3K4me3 was found to be increased in both Meg3-CAS and Meg3-TSS myoblasts compared to pX controls suggesting a widespread increase in transcription in these mutant genomes (Fig. 7B). The most striking difference in the H3K4me3 pattern, however, was noted in myotubes. Meg3-CAS myotubes had substantially more H3K4me3 than pX control and Meg3-TSS myotubes, suggesting heightened transcription activation in myotubes overexpressing *Dlk1-Dio3* ncRNAs (Fig. 7B).

The parallel and sustained increase in H3K27me3 and H3K4me3 marks in Meg3-CAS myotubes raised the possibility that these mutants had more bivalent promoters, a key indicator of developmental regulatory genes poised to be rapidly activated during differentiation (Blanco et al. 2020; Klein and Hainer, 2020; Macrae et al., 2023). Therefore, we analyzed both mutant genomes for potential overlap in these histone marks to determine whether bivalent promoters were more prevalent in Meg3-CAS mutants. As shown in Fig. 7C, the number of bivalent regions were similar in both mutant myoblast genomes, but in myotubes the number of bivalent promoters in Meg3-CAS myotubes remained high and was greater than that in Meg3-TSS mutants. Gene ontology (GO) analysis revealed that the bivalent promoters in both mutant myoblasts were associated with cellular processes related to growth, differentiation, development, and morphogenesis (Fig. 7D).

Interestingly, GO analysis of bivalent marks in myotubes showed that Meg3-CAS mutants continued to have substantially more poised promoters associated with developmental or differentiation pathways compared to Meg3-TSS and pX control myotubes (Fig. 7E). Taken together, these results demonstrate not only the differential sensitivity of histone methylation modifications to *Dlk1-Dio3* ncRNA dosage but also how their overexpression has a more impactful modulatory effect on poised promoters in muscle cells.

## Discussion

Of the multiple steps needed for proper muscle differentiation, e.g. extracellular signals, cell-cell contact, cell cycle exit, and cell fusion, remodeling the myoblast genome to establish cell state-specific gene expression patterns during this dynamic process is among the most critical. As myoblasts transition from proliferation to differentiation there are also major changes in metabolic demands. Thus, metabolism and genome remodeling must be tightly controlled to activate the proper repertoire of gene programs for efficient differentiation and prevent an abnormal cell state. Here we have shown that the *Dlk1-Dio3* ncRNA cluster is a key modulator of muscle cell state through its coordinate regulation of mitochondrial respiration and histone methylation. Dissecting their collective regulatory function was achieved through carefully designed CRISPR/Cas9-mediated deletions in the *Meg3* proximal promoter region which resulted in misexpression of the entire polycistronic ncRNA transcript. Our data demonstrates that the *Dlk1-Dio3* ncRNA cluster is an essential component in a metabolo-epigenomic network necessary for regulating cell state in the myogenic lineage. Changes in its dosage inappropriately activated muscle sarcomere genes and mitochondrial respiration (OXPHOS) and reprogrammed chromatin structure in myoblasts to severely impede differentiation.

Approaches to define the combined function of the *Dlk1-Dio3* ncRNA cluster have been hampered by its large and unusually elaborate gene structure. This mammalian-specific ncRNA locus has evolved to encode three different classes of ncRNAs, each harboring multiple copies, that are transcribed simultaneously. To date, studies have largely focused on investigating portions of the gene including deletions of selected miRNA sub-clusters, or misexpression of its individual ncRNAs, which have resulted in overlapping but largely distinct phenotypes (Kameswaran et al., 2014; Labialle et al., 2014; Qian et al., 2016; Vu Hong et al., 2022; Wust et al., 2018). Our results differ in two key respects from prior studies. First, we altered the dosage of the entire *Dlk1-Dio3* ncRNA cluster, and not a subset, through specific deletions of *cis*-regulatory sequences in the *Meg3* proximal promoter region in myoblasts. Remarkably, one of these proximal promoter deletions caused a dramatic and unexpected increase in expression of the >200kb *Dlk1-Dio3* ncRNA cluster in its native genomic context, suggesting the removal of a repressive *cis*-acting sequence(s). Second, unlike the IG-DMR mutations, dysregulated expression induced by both promoter deletions was restricted to the ncRNA cluster and did not have any significant far-reaching effects on the *Dlk1* and *Dio3* protein-coding genes, providing a more direct and comprehensive analysis of *Dlk1-Dio3* ncRNA function.

Misexpression of the *Dlk1-Dio3* ncRNA cluster resulted in impaired myotube formation which was accompanied by aberrant expression of sarcomeric and OXPHOS proteins and downregulation of EMT transcripts, suggesting a disruption in cell state and highlighting the importance of dosage in maintaining a stable myogenic identity, consistent with the biology of developmentally imprinted genes. Interestingly, although transcriptomic and pathway analyses predicted significant enrichment in OXPHOS and myogenesis in both mutant lines, these mutants were functionally distinct. For example, the upregulation of OXPHOS transcripts in both mutant lines resulted in dramatically different mitochondrial respiration and membrane potential. While a small fraction of preferentially dysregulated transcripts could explain these functional differences, it is conceivable that differences in respiration relate to the specific complement or function of OXPHOS transcripts in each gene set, e.g. more positive-acting OXPHOS factors in the Meg3-CAS upregulated gene set compared to Meg3-TSS. Similarly, myogenesis related transcripts were upregulated in both mutant lines but their myotube morphologies were noticeably different, possibly due to preferential dysregulation in the types of structural proteins expressed or their post-translational modifications. Nevertheless, these results indicate that differentiation and mitochondrial metabolism were uniquely affected by contrasting dosage of *Dlk1-Dio3* ncRNAs.

We also sought to reconcile how ETC protein expression patterns were driving functional differences in mitochondrial respiration between the two mutant lines. The reduction in OCR in Meg3-TSS myoblasts could be related to the decreased expression of SDHB affecting two distinct metabolic pathways in mitochondria. As SDHB catalyzes reactions in the TCA cycle and ETC, its downregulation could impair both Complex II activity (ETC) and the TCA cycle. In the TCA cycle, reduced oxidation of succinate to fumarate would generate fewer electrons transferred to the cofactor FAD to generate FADH_2_. This could impair OXPHOS by having not only fewer molecules of FADH_2_ available at Complex II but also decreasing the flow of electrons from FADH_2_ into the intermembrane space - the other function of SDHB - resulting in reduced OCR and ATP production. Regarding Meg3-CAS myoblasts, Seahorse analysis detected significantly increased proton leak in these mutants, consistent with a reduction in MitoTracker CMXRos signal, but contrary to the observed increase in respiration. Intriguingly, the increase in MT-CO1 (Complex IV) expression in Meg3-CAS myoblasts and myotubes could mitigate increased protein leak by driving respiration and maintaining ATP production despite the inefficiency introduced by the leak. Taken together, the distinct expression profile of ETC protein subunits helps explain the differences in mitochondrial respiration detected in the mutants. Moreover, perturbations in the timing of OXPHOS activation in mutant myoblasts could contribute to defective muscle differentiation given the importance of coordinating the appropriate energy-yielding reaction(s) with a specific cell state in myogenesis.

While *Meg3* has been established to interact with PRC2 to mediate the repressive H3K27me3 mark, the effects of the whole *Dlk1-Dio3* ncRNA cluster had not previously been linked to epigenetic regulation. We observed through ATAC-seq analysis genome-wide changes in chromatin accessibility in both Meg3-CAS and Meg3-TSS. Contrary to the upregulated expression of OXPHOS transcripts found by RNA-seq, OXPHOS loci were less accessible in Meg3-CAS and Meg3-TSS myoblasts and myotubes compared to pX controls. Though counterintuitive this may reflect differences in regulatory complexity, i.e. the number of accessible regions in the proximity of a gene. It has been described that changes in gene expression do not always correspond to increased accessibility for low complexity genes (Gonzalez et al., 2015; Kiani et al., 2022). Even though OXPHOS transcripts were upregulated in *Dlk1-Dio3* ncRNA mutants, chromatin accessibility at these loci might not change to a great extent particularly if chromatin in these regions already exists in a relatively open conformation in wild type myoblasts. Rather upregulated expression could relate to increased transcription factor activity already bound at these regulatory regions or displacement of repressive factors resulting in enhanced transcription without requiring more accessibility.

Differences in the extent of histone methylation marks H3K27me3 and H3K4me3 was also evident between the two mutants. Overexpression of *Dlk1-Dio3* ncRNAs resulted in more H3K27me3 and H3K4me3 in myotubes leading to a greater number of bivalent promoters in the Meg3-CAS myotube genome. While the molecular mechanisms establishing bivalent promoters are complex, PRC2 and the Trithorax family member, KMT2B, are considered the predominant histone methyltransferases for histone modifications in these regions (Blanco et al., 2020; Klein and Hainer, 2020; Macrae et al., 2023). The increased H3K27me3 in bivalent promoters in Meg3-CAS myotubes is entirely consistent with the increased abundance of *Dlk1-Dio3* lncRNAs such as *Meg3* contributing to greater PRC2 recruitment at these regions. Although less is known about lncRNA-mediated recruitment of KMTs, the recruitment of this class of histone methyltransferases by the *Dlk1-Dio3* ncRNA cluster warrants further investigation.

The observed defects in *Dlk1-Dio3* ncRNA mutants presumably stem from a complex interplay of the various classes of mature ncRNAs which function in distinct subcellular compartments (nucleus, nucleolus, cytoplasm) and participate in disparate cell regulatory processes ranging from chromatin structure (lncRNAs) to translation (miRNAs and snoRNAs). Although nothing is known about the post-transcriptional processing mechanisms and regulation of the polycistronic primary RNA, another layer of complexity arises from the synergistic and antagonistic interactions of these fully processed ncRNAs and their respective targets. Lastly, as we have shown in this study, expression timing and dosage of *Dlk1-Dio3* ncRNAs are important determinants in the regulation of muscle cell state and differentiation as summarized in the model depicted in Fig. 8. But some outstanding questions remain. Pathway analyses revealed significant dysregulation of other mitochondrial-related processes such as mitophagy and reactive oxygen species (ROS). Are these processes differentially affected? And, if so, does this help explain functional differences in proton leak and membrane potential in these mutants? In addition, investigating the discordance between chromatin structure and transcriptional output in these mutants will require more extensive analysis of regulatory regions and DNA motifs at dysregulated loci. Our study has highlighted the interconnectedness of mitochondrial metabolic activity and the epigenome and how disruption of this finely tuned coregulation by the *Dlk1-Dio3* ncRNA cluster radically alters myogenic cell state. This information can be used to further understand how breakdown in metabolic and epigenetic regulation contribute to degeneration and inefficient myogenesis and regeneration observed in dystrophic muscle.

**Fig. 8.**
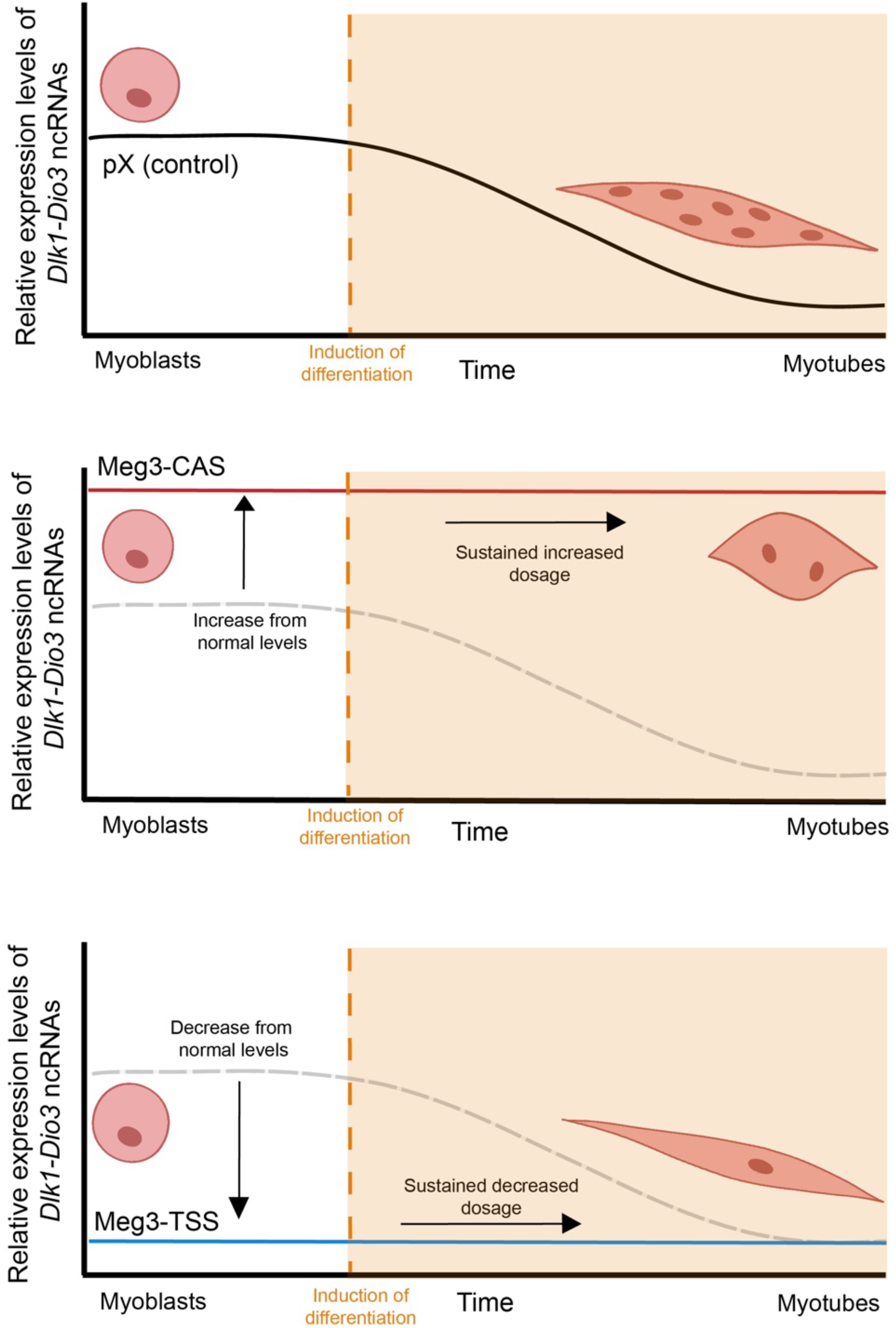
Model depicting importance of dosage and timing of *Dlk1-Dio3* ncRNA expression for myogenesis. (Top panel) Efficient skeletal muscle differentiation requires the correct dosage of *Dlk1-Dio3* ncRNAs at the proper time which has been shown to be critical in regulating metabolism, cell state transitions, and EMT. (Middle panel) Sustained upregulation of *Dlk1-Dio3* ncRNAs throughout the differentiation process results in impaired myotube formation. (Bottom panel) Sustained downregulation of *Dlk1-Dio3* ncRNAs throughout the differentiation process results in impaired myotube formation.

## MATERIALS AND METHODS

### CRISPR-Cas9 Plasmid Cloning

The pX458 pSpCas9 mammalian expression vector was provided by the Pajevic Lab (Boston University School of Medicine). The pX458 pSpCas9 construct was digested with BbsI restriction enzyme and the linearized vector extracted. Pairs of guide RNAs were designed to target regions of the *Meg3* proximal regulatory region for deletion using *in silico* design tools provided through MIT (https://crispr.mit.edu) and the Broad Institute (https://portals.broadinstitute.org/gpp/public/analysis-tools/sgrna-design). DNA oligonucleotides corresponding to each guide RNA sequence (Table S1) were cloned into the pX458 pSpCas9 mammalian expression vector.

### C2C12 Cell Culture and Transfection

C2C12 myoblast cells were cultured in growth medium consisting of Dulbecco’s Modified Eagle Medium (DMEM) supplemented with 10% fetal bovine serum (FBS), and 1% 100X penicillin-streptomycin L-glutamine (P/S/G). Myoblasts were transfected with PEI and two of the *Meg3* promoter-specific CRISPR-Cas9/gRNA constructs or the pX458 vector alone and 72 h later EGFP positive cells selected by FACS. A population of 3000 - 10000 EGFP positive cells was collected at the end of each sort and serially diluted to seed one GFP-positive per well onto 96-well plates. A population of pX458 transfected cells was obtained for use as a control in experiments. Prior to differentiation, C2C12 myoblasts were grown in growth medium to confluence. To induce differentiation, the media was changed to differentiation media consisting of DMEM, 2% horse serum, and 1% P/S/G.

### PCR Genotyping Validation of Successful CRISPR-Cas9 Deletion

DNA was isolated from the CRISPR-edited clonal C2C12 cells and PCR genotyping was then used to confirm the presence of CRISPR-Cas9 deletions by using the primers flanking the region of the *Meg3* promoter that was targeted for deletion (Table S2). Following confirmation of homozygous deletion, the gel purified PCR product was confirmed by sequencing. The two clones with the desired genomic edits were used for subsequent experiments.

### cDNA Synthesis and qPCR

RNA was isolated from C2C12 cells using TRIzol reagent (Invitrogen) followed by phenol-chloroform extraction, precipitated with 100% isopropanol, and washed with 70% ethanol. RNA pellets were air-dried and resuspended in RNase-free water. RNA template was converted to cDNA using M-MLV-Reverse Transcriptase (Promega) according to the manufacturer’s protocol. Quantitative RT-PCR was performed in triplicate wells using Power SYBR Green Master Mix (Applied Biosystems) with the 7900HT Sequence Detection System (Applied Biosystems). Primers used for qRT-PCR analyses are listed in Table S3.

### Immunocytochemistry

pX, Meg3-CAS, and Meg3-TSS myoblasts were seeded at a density of 25,000 cells and allowed to grow for 24 h for myoblast experiments and 100,000 cells and allowed to grow for 48 h for myotube experiments, on a glass coverslip pre-treated with 0.1% gelatin. To investigate cleaved Caspase-3, myoblasts were grown to confluence and subjected to differentiation media for 12 h before beginning immunostaining to represent day 0 of differentiation. For myotube experiments, after 48 h the media was switched to differentiation media, and the cells were then differentiated as previously described for three days. The myoblasts or myotubes were then fixed with 4% paraformaldehyde and blocked with PBS solution containing 10% donkey serum and 0.1% Triton-X. Cells were incubated with their respective primary antibody solution: α-actinin (1:1000, Sigma A7811), Cleaved Caspase-3 (Asp175) (1:200, Invitrogen PA5-114687), Tom20 (1:400, CST 42406) overnight at 4°C. The coverslips were then thoroughly washed with 1X PBS and the secondary antibody solution: donkey anti-mouse Alexa-Fluor 488 IgG (1:1000, Invitrogen A21202), donkey anti-rabbit Alexa-Fluor 555 IgG (1:1000, Invitrogen A31572), donkey anti-rabbit Alexa-Fluor 568 IgG (1:1000, Invitrogen A10042) was added for 1 h. The coverslips were thoroughly washed with 1X PBS and then mounted on microscope slides using DAPI Vectashield mounting medium. To measure mitochondrial membrane potential, myoblasts adhered to gelatin-treated coverslips were first incubated with 50nM of MitoTracker Red CMX-Ros probe (Invitrogen) for 30 minutes prior to immunostaining. Immunofluorescence images were captured for 4 fields of view per coverslip using a Nikon Eclipse (Nikon Eclipse Ni-E Motorized Fluorescent Microscope) with uniform exposure settings across all samples for quantification using FIJI. The fusion index is calculated based on the percentage of nuclei within cells, delineated by α-actinin expression, containing one nucleus, two nuclei, and more than three nuclei, with each category representing a proportion of the total nuclei within the cell border, and these percentages sum up to 100%.

### Western Blot

Whole cell lysates or isolated mitochondria were obtained from myoblasts or myotubes at day 3 of differentiation using ELB buffer or Abcam Mitochondrial Isolation Kit for Cultured Cells (ab110170), respectively. For either procedure, cells were dounce homogenized on ice, and protein concentration was determined using the Pierce BCA Protein Assay Kit (ThermoFisher). To detect mitochondrial protein expression, 30 ug of isolated mitochondria were loaded into 4-20% precast polyacrylamide gel (BioRad), electrophoresed, and then transferred to PVDF membrane using a high pH CAPS buffer (10 mM CAPS, 10% methanol, pH 11 adjusted with NaOH) for 1.5 hours at 200 mA. Membranes were blocked in 5% nonfat dry milk and then subjected to primary antibody incubation overnight at 4℃. Subsequently, membranes were washed with 1X TBS/T (0.1% Tween 20) and secondary antibodies were incubated at room temperature for 1 hour. Membranes were washed with 1X TBS/T (0.1% Tween 20) followed by a final wash with 1X TBS and dried in whatman paper prior to imaging on the Azure Sapphire. To detect myosin heavy chain expression, 50 ug of total protein lysate was used. The same procedure described above was used but the transfer buffer was modified (20 mM Tris, 150 mM glycine, 20% methanol, 0.05% SDS). Primary antibodies against the following proteins were used: ɑ-tubulin (1:1000, Cell Signaling Technology mAb #3873), MF20 (0.2 μg/ml, DSHB MF-20 supernatant), and OXPHOS (1:250, Abcam ab110413). Secondary dye used was mouse IRDye 680RD (1:2000, Licor 926-68070). The membrane was imaged on the Azure Sapphire Biomolecular Imager followed by Coomassie staining of the membrane to determine total protein content and imaged on Azure c200.

### Seahorse XF Pro Analyzer

Mitochondrial respirometry was performed using the Seahorse XF Pro Analyzer (Agilent). Cells were grown on tissue culture plates and 24 hours before testing, cells were seeded at 12,000 cells per well into a Seahorse XFe96/XF Pro Cell Culture Microplate in 80 µl of growth media. The day prior to testing, the cartridge was hydrated according to the manufacturer’s instructions. On the day of the experiment, Seahorse XF DMEM was supplemented with 1M glucose and 200 mM glutamine and warmed to 37 °C. Cells were washed with warmed assay medium and placed in a 37 °C non-CO_2_ incubator for 45 minutes prior to the assay. Immediately before starting the assay, cells were given fresh assay medium. The Seahorse XF Cell Mito Stress Test was carried out according to manufacturer instructions (Agilent 103015-100). Following the test, cells were processed in ELB buffer, and protein concentration was determined for normalization of the data to protein content per well.

### RNA-Sequencing

RNA samples corresponding to myoblasts and myotubes (day 3 of differentiation) were submitted to the Boston University Microarray and Sequencing Resource Core in triplicate for library prep and Illumina sequencing. FASTQ files were aligned to mouse genome build mm10 using STAR v2.7.9a. Ensembl-Gene-level counts were generated separately for mitochondrial and non-mitochondrial genes using featureCounts (Subread package, version 1.6.2) and Ensembl annotation build 100 (uniquely aligned proper pairs, same strand). The DESeq2 R package (v1.23.10) was used to compute variance-stabilizing transformed (VST) values and to perform Wald tests with R v4.1.2. Correction for multiple hypothesis testing was accomplished using the Benjamini-Hochberg false discovery rate (FDR). All human Entrez Gene identifiers matching exactly one mouse Ensembl Gene identifier (determined using Ensembl Biomart and NCBI ‘gene_orthologs’ table retrieved 2024-02-22) were ranked by the corresponding Wald statistic computed for each pairwise comparison. Each ranked list was then used to perform pre-ranked Gene Set Enrichment Analysis (GSEA) analyses (v2.2.1, default parameters with random seed 1234) using the Entrez Gene version of the Hallmark gene sets obtained from the Molecular Signatures Database (MSigDB), version 7.5.1.

### ATAC-Sequencing (ATAC-seq) and CUT & RUN

For ATAC-seq, Myoblast and myotube samples in duplicate or triplicate were processed according to manufacturer’s instructions (ATAC-Seq Kit Active Motif Kit 53150). For CUT&RUN, myoblast and myotube samples in triplicate were processed according to manufacturer’s instructions (Epicypher CUTANA CUT&RUN Kit (14-1048) and CUTANA CUT&RUN Library Prep Kit (14-1001)). DNA libraries were sequenced using the Illumina NextSeq 2000 to obtain 30-45 million reads per sample. Sequenced DNA libraries were then quality-checked using FastQC (Andrews, 2010). For CUT&RUN, reads were trimmed for adapters and reads under 25 base pairs in length were removed using Trimmomatic v0.36 (Bolger et al., 2014). Reads for both libraries were aligned to the *Mus musculus* reference genome mm10 using Bowtie 2 v2.4.2 (Langmead and Salzberg, 2012). PCR duplicates were identified using Picard’s MarkDuplicates v2.8.0 (Broad Institute, 2019) and filtered using SAMtools v1.12 (Danecek et al., 2021). For CUT&RUN, the spike-in was aligned to the *E. coli* genome. For ATAC-seq, reads aligning to blacklist regions specified by ENCODE were removed using BEDtools v2.31.0 (Quinlan and Hall, 2010; Amemiya et al., 2019). ATAC-seq peaks were called using MACS2 v2.2.7.1 with default parameters for paired-end analysis (-f BAMPE) and a q-value cutoff of 0.05 (Zhang et al., 2008). For CUT&RUN, peaks were called using SEACR v1.3 using the IgG control track previously normalized to *E. coli* with stringent threshold (non stringent) (Meers et al., 2019). Identified peaks and read density were visualized using IGV (Robinson et al., 2011) alongside the ENCODE cis-regulatory elements track (https://genome.ucsc.edu/cgi-bin/hgTrackUi?db=mm10&g=encodeCcreCombined).

Differentially expressed peaks were identified using the edgeR differential model in the DiffBind v3.14.0 package (Stark and Brown, 2011). Subsequent annotation of these differential peaks to their closest genes was performed using Homer v4.11 (Heinz et al., 2010).

### Statistical Analysis

Numerical quantification is represented as the mean ±s.e.m. of at least three replicates. Statistical significance was determined using one-way ANOVA with *p* < 0.05 being the threshold for significance, followed by Tukey’s multiple comparisons test. (*p<0.05, **p<0.01, ***p<0.001, ****p<0.0001).

## Acknowledgments

We are grateful to the BU Microarray and Sequencing Resource for RNA sequencing, ATAC sequencing, and CUT&RUN sequencing. We thank Mahekpreet Pannu, Eliza Miller, Kaina Millan, Anna Mcniff for their technical assistance.

## Data availability

The RNA-seq data have been deposited in NCBI’s Gene Expression Omnibus and are accessible through GEO Series accession number GSE268482 (https://www.ncbi.nlm.nih.gov/geo/query/acc.cgi?acc=GSE268482).

---

**Figure S1:**
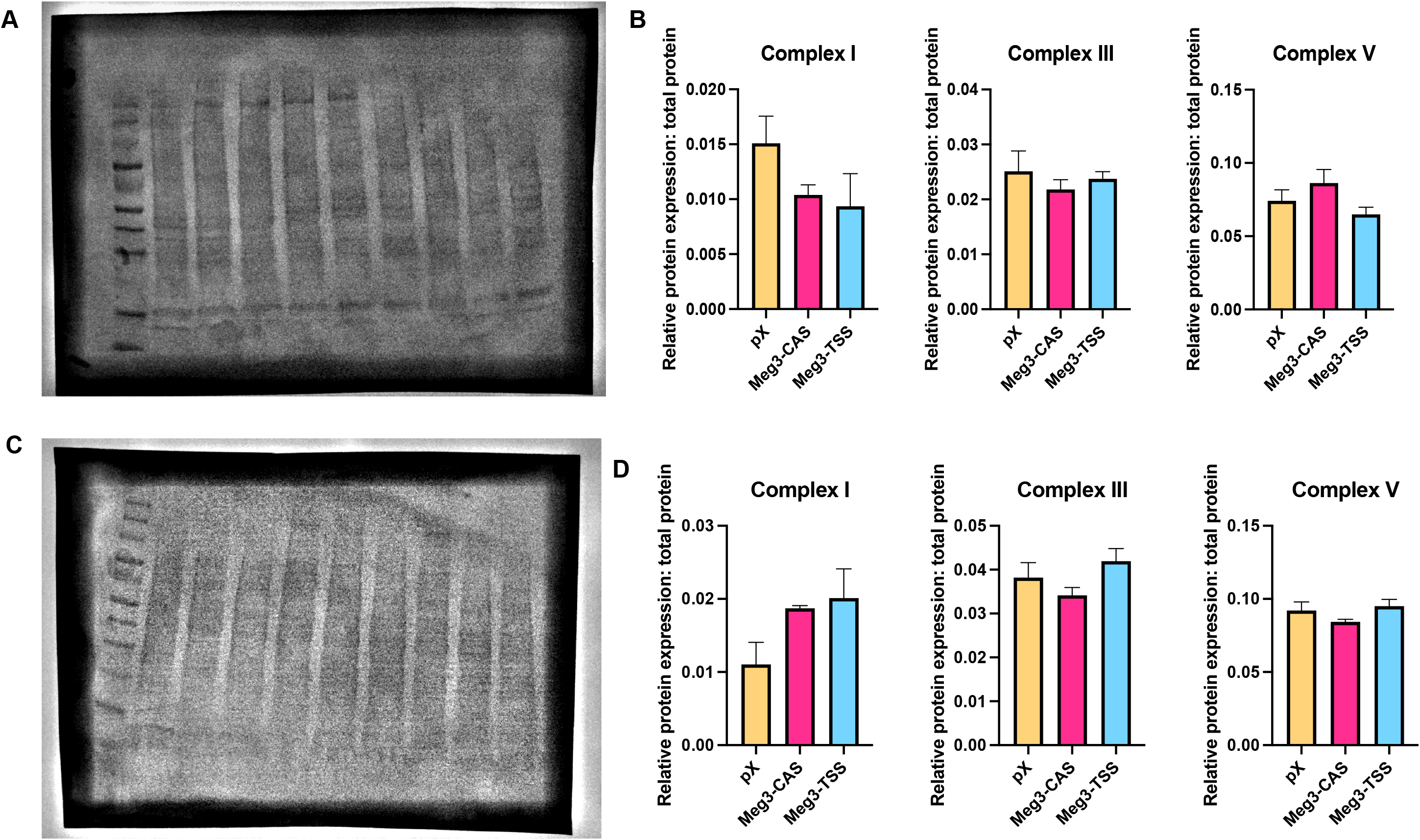
Protein expression levels of select OXPHOS complex proteins remain unchanged and myogenic genes are differentially expressed in *Dlk1-Dio3* ncRNA mutant cell lines. A) Coomassie on PVDF membrane for total protein from isolated mitochondria from myoblasts used to investigate OXPHOS proteins. B) Complex I, Complex III, and Complex V have no difference in protein expression across pX, Meg3-CAS, and Meg3-TSS myoblasts cells. C) Coomassie on PVDF membrane for total protein from isolated mitochondria from myotubes used to investigate OXPHOS proteins. D) Complex I, Complex III, and Complex V have no difference in protein expression across pX, Meg3-CAS, and Meg3-TSS myotubes cells.

**Figure S2:**
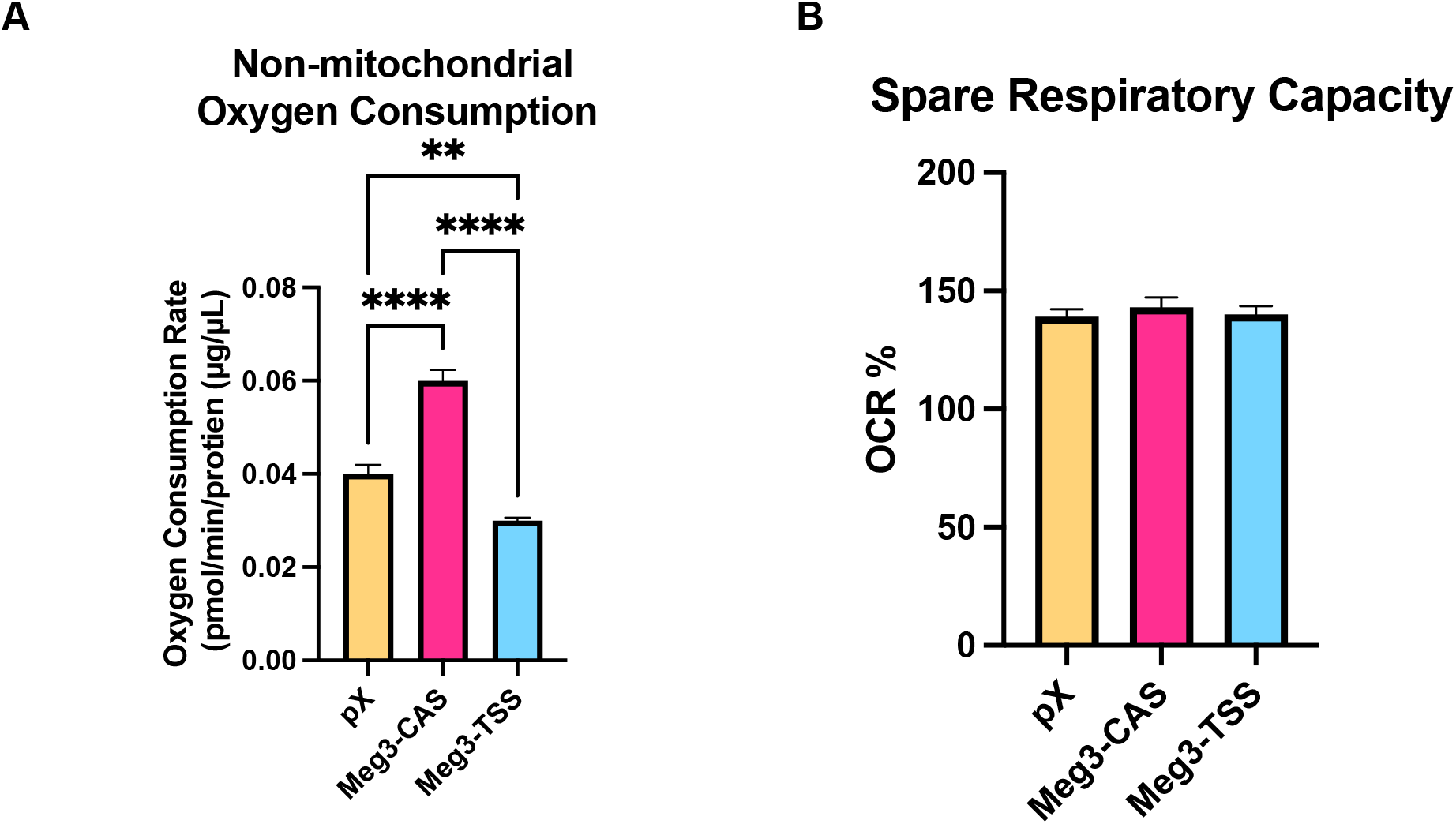
Mitochondrial respiration measurements show differential non-mitochondrial oxygen consumption in mutant myoblasts but no difference in spare respiratory capacity as a percentage of OCR. A) Non-mitochondrial oxygen consumption was increased in Meg3-CAS myoblasts and decreased in Meg3-TSS myoblasts compared to pX myoblasts. B) Spare respiratory capacity as a percentage of OCR was unchanged across the cell lines at the myoblasts stage.

**Figure S3:**
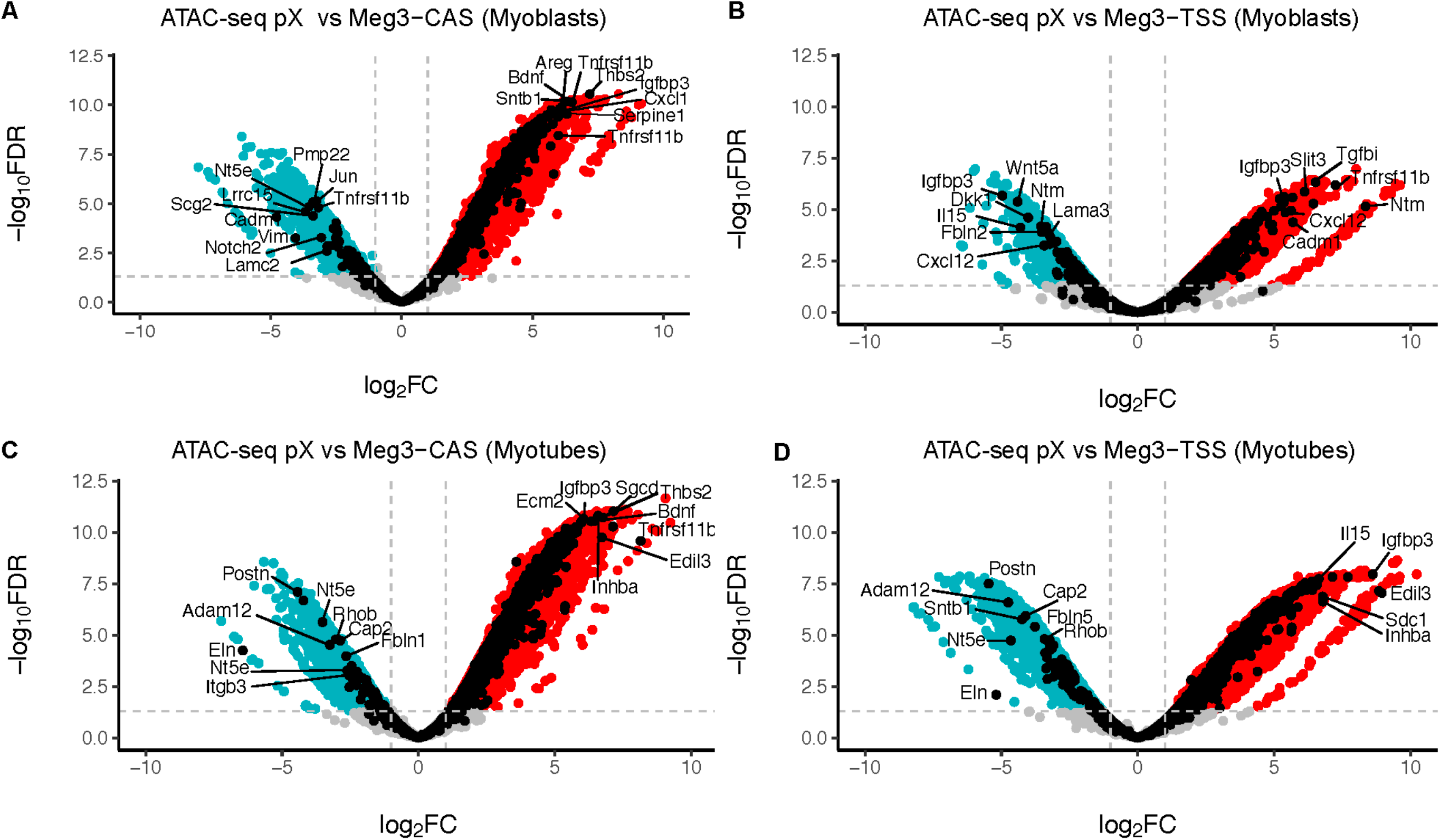
Differential chromatin landscape for epithelial-mesenchymal transition genes in cells with altered *Dlk1-Dio3* ncRNA expression. (A) Volcano plot of ATAC-seq data comparing differentially open regions of chromatin in pX compared to Meg3-CAS myoblasts, with EMT genes denoted in black (Top 10 unique EMT genes labeled). (B) Volcano plot of ATAC-seq data comparing differentially open regions of chromatin in pX compared to Meg3-TSS myoblasts, with EMT genes denoted in black (Top 10 unique EMT genes labeled). (C) Volcano plot of ATAC-seq data comparing differentially open regions of chromatin in pX compared to Meg3-CAS myotubes, with EMT genes denoted in black (Top 10 unique EMT genes labeled). (D) Volcano plot of ATAC-seq data comparing differentially open regions of chromatin in pX compared to Meg3-TSS myotubes, with EMT genes denoted in black (Top 10 unique EMT genes labeled).

## References

Amemiya, H.M., Kundaje, A. and Boyle, A.P. (2019). ‘The ENCODE Blacklist: Identification of Problematic Regions of the Genome’. Scientific Reports. 9(1), p. 9354.

Andrews, S. (2010). FastQC: A quality control tool for high throughput sequence data. Available at: https://www.bioinformatics.babraham.ac.uk/projects/trim_galore/.

Benetatos, L., Hatzimichael, E., Londin, E., Vartholomatos, G., Loher, P., Rigoutsos, I., and Briasoulis, E. (2013). The microRNAs within the DLK1-DIO3 genomic region: involvement in disease pathogenesis. Cell Mol Life Sci. 70(5), 795–814.

Bhattacharya, D., and Scimè, A. (2020). Mitochondrial Function in Muscle Stem Cell Fates. Front Cell Dev Biol. 8, 480.

Blanco E., González-Ramírez M., Alcaine-Colet A., Aranda S., Di Croce L. (2020). The Bivalent Genome: Characterization, Structure, and Regulation. Trends Genet. 36(2), 118–131.

Bolger, A.M., Lohse, M. and Usadel, B. (2014). ‘Trimmomatic: a flexible trimmer for Illumina sequence data’. Bioinformatics. 30(15), 2114–2120.

Bourdeau Julien, I., Sephton, C.F., and Dutchak, P.A. (2018). Metabolic Networks Influencing Skeletal Muscle Fiber Composition. Front Cell Dev Biol. 6, 125.

Broad Institute. (2019). ‘Picard Toolkit’. Available at: https://broadinstitute.github.io/picard/.

Chakrabarty, R.P., and Chandel, N.S. (2021). Mitochondria as Signaling Organelles Control Mammalian Stem Cell Fate. Cell Stem Cell. 28(3), 394–408.

Danecek, P. et al. (2021). ‘Twelve years of SAMtools and BCFtools’. GigaScience. 10(2), giab008.

da Rocha, S.T., Edwards, C.A., Ito, M., Ogata, T., and Ferguson-Smith, A.C. (2008). Genomic imprinting at the mammalian Dlk1-Dio3 domain. Trends Genet. 24(6), 306–16.

Das, S., Morvan, F,. Morozzi, G., Jourde, B., Minetti, G.C., Kahle, P., Rivet, H., Brebbia, P., Toussaint, G., Glass, D.J., et al. (2017). ATP Citrate Lyase Regulates Myofiber Differentiation and Increases Regeneration by Altering Histone Acetylation. Cell Rep. 21(11), 3003–3011.

Dill, T.L., and Naya, F.J. (2018). A Hearty Dose of Noncoding RNAs: The Imprinted DLK1-DIO3 Locus in Cardiac Development and Disease. J Cardiovasc Dev Dis. 5(3).

Dill, T.L., Carroll, A., Pinheiro, A., Gao, J., and Naya, F.J. (2021). The long noncoding RNA Meg3 regulates myoblast plasticity and muscle regeneration through epithelial-mesenchymal transition. Development. 148(2), dev194027.

Edwards, C.A., Mungall, A.J., Matthews, L., Ryder, E., Gray, D.J., Pask, A.J., Shaw, G., Graves, J.A., Rogers, J.; SAVOIR consortium; et al. (2008). The evolution of the DLK1-DIO3 imprinted domain in mammals. PLoS Biol. 6(6), e135.

Egan, B., and Zierath, J.R. (2013). Exercise metabolism and the molecular regulation of skeletal muscle adaptation. Cell Metab. 17(2), 162–84.

Etchegaray, J.P., and Mostoslavsky, R. (2016). Interplay between Metabolism and Epigenetics: A Nuclear Adaptation to Environmental Changes. Mol Cell. 62(5), 695–711.

Folmes, C.D., Dzeja, P.P., Nelson, T.J., and Terzic, A. (2012). Metabolic plasticity in stem cell homeostasis and differentiation. Cell Stem Cell. 11(5), 596–606.

Fontes-Oliveira, C.C., Steinz, M., Schneiderat, P., Mulder, H., and Durbeej, M. (2017). Bioenergetic Impairment in Congenital Muscular Dystrophy Type 1A and Leigh Syndrome Muscle Cells. Sci Rep. 7, 45272.

Gao, Y.Q., Chen, X., Wang, P., Lu, L., Zhao, W., Chen, C., Chen, C.P., Tao, T., Sun, J., Zheng, Y.Y., et al. (2015). Regulation of DLK1 by the maternally expressed miR-379/miR-544 cluster may underlie callipyge polar overdominance inheritance. Proc Natl Acad Sci U S A. 112(44), 13627–32.

González, A.J., Setty, M., and Leslie, C.S. (2015). Early enhancer establishment and regulatory locus complexity shape transcriptional programs in hematopoietic differentiation. Nat Genet. 47(11), 1249–59.

Groh, S., Zong, H., Goddeeris, M. M., Lebakken, C. S., Venzke, D., Pessin, J. E., and Campbell, K. P. (2009). Sarcoglycan complex: implications for metabolic defects in muscular dystrophies. J. Biol. Chem. 284, 19178–19182

Hardee, J.P., Martins, K.J.B., Miotto, P.M, Ryall, J.G., Gehrig, S.M., Reljic, B., Naim, T., Chung, J.D., Trieu, J., Swiderski, K., et al. (2021). Metabolic remodeling of dystrophic skeletal muscle reveals biological roles for dystrophin and utrophin in adaptation and plasticity. Mol Metab. 45, 101157.

Hargreaves, M., and Spriet, L.L. (2020). Skeletal muscle energy metabolism during exercise. Nat Metab. 2(9), 817–828.

Heinz, S. et al. (2010). ‘Simple Combinations of Lineage-Determining Transcription Factors Prime cis-Regulatory Elements Required for Macrophage and B Cell Identities’. Mol Cell. 38(4), 576–589.

Heydemann, A. (2018). Skeletal Muscle Metabolism in Duchenne and Becker Muscular Dystrophy-Implications for Therapies. Nutrients. 10(6), 796.

Kameswaran, V., Bramswig, N.C., McKenna, L.B., Penn, M., Schug, J., Hand, N.J., Chen, Y., Choi, I., Vourekas, A., Won, K.J., et al. (2014). Epigenetic regulation of the DLK1-MEG3 microRNA cluster in human type 2 diabetic islets. Cell Metab. 19(1), 135–45.

Kaneko, S., Bonasio, R., Saldaña-Meyer, R., Yoshida, T., Son, J., Nishino, K., Umezawa, A., and Reinberg, D. (2014). Interactions between JARID2 and noncoding RNAs regulate PRC2 recruitment to chromatin. Mol Cell. 53(2), 290–300.

Kiani, K., Sanford, E.M., Goyal, Y., and Raj, A. (2022). Changes in chromatin accessibility are not concordant with transcriptional changes for single-factor perturbations. Mol Syst Biol. 18(9), e10979.

Kircher, M., Bock, C., and Paulsen, M. (2008). Structural conservation versus functional divergence of maternally expressed microRNAs in the Dlk1/Gtl2 imprinting region. BMC Genomics. 9, 346.

Klein D.C. and Hainer S.J. (2020). Chromatin regulation and dynamics in stem cells. Curr Top Dev Biol. 138, 1–71.

Labialle, S., Marty, V., Bortolin-Cavaillé, M.L. Hoareau-Osman, M., Pradère, J.P., Valet, P., Martin, P.G., and Cavaillé, J. (2014). The miR-379/miR-410 cluster at the imprinted Dlk1-Dio3 domain controls neonatal metabolic adaptation. EMBO J. 33(19), 2216–30.

Langmead, B. and Salzberg, S.L. (2012). ‘Fast gapped-read alignment with Bowtie 2’. Nature Methods, 9(4), 357–359.

Lin, S.P., Youngson, N., Takada, S., Seitz, H., Reik, W., Paulsen, M., Cavaille, J. and Ferguson-Smith, A.C. (2003). Asymmetric regulation of imprinting on the maternal and paternal chromosomes at the Dlk1-Gtl2 imprinted cluster on mouse chromosome 12. Nature genetics. 35(1), 97–102.

Ly, C.H., Lynch, G.S., and Ryall, J.G. (2020). A Metabolic Roadmap for Somatic Stem Cell Fate. Cell Metab. 31(6), 1052–1067.

Macrae, T.A., Fothergill-Robinson, J., and Ramalho-Santos, M. (2023). Regulation, functions and transmission of bivalent chromatin during mammalian development. Nat Rev Mol Cell Biol. 24(1), 6–26.

McGee, S.L., Fairlie, E., Garnham, A.P., and Hargreaves, M. (2009). Exercise-induced histone modifications in human skeletal muscle. J Physiol. 587(Pt 24), 5951–8.

Meers, M.P., Tenenbaum, D. and Henikoff, S. (2019). Peak calling by Sparse Enrichment Analysis for CUT&RUN chromatin profiling. Epigenetics & Chromatin. 12(1), 42.

Nguyen, J.H., Chung, J.D., Lynch, G.S., and Ryall, J.G. (2019). The Microenvironment Is a Critical Regulator of Muscle Stem Cell Activation and Proliferation. Front Cell Dev Biol. 7, 254.

Nieborak, A., and Schneider, R. (2018). Metabolic intermediates - Cellular messengers talking to chromatin modifiers. Mol Metab. 14, 39–52.

Pillon, N.J., Sardón Puig, L., Altıntaş, A., Kamble, P.G., Casaní-Galdón, S., Gabriel, B.M., Barrès, R., Conesa, A., Chibalin, A.V., Näslund, E., et al. (2022). Palmitate impairs circadian transcriptomics in muscle cells through histone modification of enhancers. Life Sci Alliance. 6(1), e202201598.

Qian, P., He, X.C., Paulson, A., Li, Z., Tao, F., Perry, J.M., Guo, F., Zhao, M., Zhi, L., Venkatraman, A., et al. (2016). The Dlk1-Gtl2 Locus Preserves LT-HSC Function by Inhibiting the PI3K-mTOR Pathway to Restrict Mitochondrial Metabolism. Cell Stem Cell. 18(2), 214–28.

Quinlan, A.R. and Hall, I.M. (2010). BEDTools: a flexible suite of utilities for comparing genomic features. Bioinformatics. 26(6), 841–842.

Reid, M.A., Dai, Z., and Locasale, J.W. (2017). The impact of cellular metabolism on chromatin dynamics and epigenetics. Nat Cell Biol. 19(11), 1298–1306.

Relaix, F., Bencze, M., Borok, M.J., Der Vartanian, A., Gattazzo, F., Mademtzoglou, D., Perez-Diaz, S., Prola, A., Reyes-Fernandez, P.C., Rotini, A., et al. (2021). Perspectives on skeletal muscle stem cells. Nat Commun. 12(1), 692.

Robinson, J.T. et al. (2011). Integrative genomics viewer. Nature Biotechnology. 29(1), 24–26.

Ryall, J.G., Cliff, T., Dalton, S., and Sartorelli, V. (2015). Metabolic Reprogramming of Stem Cell Epigenetics. Cell Stem Cell. 17(6), 651–662.

Schiaffino, S., and Reggiani, C. (2011). Fiber types in mammalian skeletal muscles. Physiol Rev. 91(4), 1447–531.

Sincennes, M.C., Brun, C.E., Lin, A.Y.T., Rosembert, T., Datzkiw, D., Saber, J., Ming, H., Kawabe, Y.I., and Rudnicki, M.A. (2021). Acetylation of PAX7 controls muscle stem cell self-renewal and differentiation potential in mice. Nat Commun. 12(1), 3253.

Snyder, C.M., Rice, A.L., Estrella, N.L., Held, A., Kandarian, S.C., and Naya, F.J. (2013). MEF2A regulates the Gtl2-Dio3 microRNA mega-cluster to modulate WNT signaling in skeletal muscle regeneration. Development. 140(1), 31–42.

Stark, R. and Brown, G (2011). DiffBind: differential binding analysis of ChIP-Seq peak data. Available at: http://bioconductor.org/packages/release/bioc/vignettes/DiffBind/inst/doc/DiffBind.pdf.

Takahashi, N., Okamoto, A., Kobayashi, R., Shirai, M., Obata, Y., Ogawa, H., Sotomaru, Y. and Kono, T. (2009). Deletion of Gtl2, imprinted non-coding RNA, with its differentially methylated region induces lethal parent-origin-dependent defects in mice. Human molecular genetics, 18(10), 1879–1888.

Tarazona, O.A., and Pourquié, O. (2020). Exploring the Influence of Cell Metabolism on Cell Fate through Protein Post-translational Modifications. Dev Cell. 54(2), 282–292.

Uroda, T., Anastasakou, E., Rossi, A., Teulon, J.M., Pellequer, J.L., Annibale, P., Pessey, O., Inga, A., Chillón, I., and Marcia, M. (2019). Conserved Pseudoknots in lncRNA MEG3 Are Essential for Stimulation of the p53 Pathway. Mol Cell. 75(5), 982–995.

Vu Hong, A., Bourg, N., Sanatine, P., Poupiot, J., Charton, K., Gicquel, E., Massourides, E., Spinazzi, M., Richard, I., and Israeli, D. (2022). Dlk1-Dio3 cluster miRNAs regulate mitochondrial functions in the dystrophic muscle in Duchenne muscular dystrophy. Life Sci Alliance. 6(1), e202201506.

Weinberg-Shukron, A., Youngson, N.A., Ferguson-Smith, A.C., and Edwards, C.A. (2023). Epigenetic control and genomic imprinting dynamics of the Dlk1-Dio3 domain. Front Cell Dev Biol. 11, 1328806.

Wüst, S., Dröse, S., Heidler, J., Wittig, I., Klockner, I., Franko, A., Bonke, E., Günther, S., Gärtner, U., Boettger, T., and Braun, T. (2018). Metabolic Maturation during Muscle Stem Cell Differentiation Is Achieved by miR-1/133a-Mediated Inhibition of the Dlk1-Dio3 Mega Gene Cluster. Cell Metab. 27(5), 1026–1039.

Yen, Y.P., Hsieh, W.F., Tsai, Y.Y., Lu, Y.L., Liau, E.S., Hsu, H.C., Chen, Y.C., Liu, T.C., Chang, M., Li, J., et al. (2018). *Dlk1-Dio3* locus-derived lncRNAs perpetuate postmitotic motor neuron cell fate and subtype identity. Elife. 7, e38080.

Yucel, N., Wang, Y.X., Mai, T., Porpiglia, E., Lund, P.J., Markov, G., Garcia, B.A., Bendall, S.C., Angelo, M., and Blau, H.M. (2019). Glucose Metabolism Drives Histone Acetylation Landscape Transitions that Dictate Muscle Stem Cell Function. Cell Rep. 27(13), 3939–3955.

Zhang, Y. et al. (2008). Model-based Analysis of ChIP-Seq (MACS). Genome Biology. 9(9), R137.

Zhao, J., Dahle, D., Zhou, Y., Zhang, X., and Klibanski, A. (2005). Hypermethylation of the promoter region is associated with the loss of MEG3 gene expression in human pituitary tumors. J Clin Endocrinol Metab. 90(4), 2179–86.

Zhao, J., Ohsumi, T.K., Kung, J.T., Ogawa, Y., Grau, D.J., Sarma, K., Song, J.J., Kingston, R.E., Borowsky, M., and Lee, J.T. (2010). Genome-wide identification of polycomb-associated RNAs by RIP-seq. Mol Cell. 40(6), 939–53.

Zhou, Y., Cheunsuchon, P., Nakayama, Y., Lawlor, M.W., Zhong, Y., Rice, K.A., Zhang, L., Zhang, X., Gordon, F.E., Lidov, H.G., et al. (2010). Activation of paternally expressed genes and perinatal death caused by deletion of the Gtl2 gene. Development. 137(16), 2643–52.

Zhu, W., Botticelli, E.M., Kery, R.E., Mao, Y., Wang, X., Yang, A., Wang, X., Zhou, J., Zhang, X., Soberman, R.J., et al. (2019). Meg3-DMR, not the Meg3 gene, regulates imprinting of the Dlk1-Dio3 locus. Dev Biol. 455(1), 10–18.

